# Anticipatory Eye Gaze as a Marker of Memory

**DOI:** 10.1101/2024.08.14.607869

**Authors:** Schmidig Jean Flavio, Yamin Daniel, Sharon Omer, Nadu Yoav, Nir Jonathan, Ranganath Charan, Nir Yuval

## Abstract

Human memory is typically studied by direct questioning, and the recollection of events is investigated through verbal reports. Thus, current research confounds memory per-se with its report. Critically, the ability to investigate memory retrieval in populations with deficient verbal ability is limited. Here, using the MEGA (Memory Episode Gaze Anticipation) paradigm, we show that monitoring anticipatory gaze using eye tracking can quantify memory retrieval without verbal report. Upon repeated viewing of movie clips, eye gaze patterns anticipating salient events can quantify their memory traces seconds before these events appear on the screen. A series of five experiments with a total of 145 participants using either tailor-made animations or naturalistic movies consistently reveal that accumulated gaze proximity to the event can index memory. Machine learning-based classification can identify whether a given viewing is associated with memory for the event based on single-trial data of gaze features. Detailed comparison to verbal reports establishes that anticipatory gaze marks recollection of associative memory about the event, whereas pupil dilation captures familiarity. Finally, anticipatory gaze reveals beneficial effects of sleep on memory retrieval without verbal report, illustrating its broad applicability across cognitive research and clinical domains.

## Introduction

Traditionally, human memory assessment has predominantly relied on explicit retrieval tasks, where participants verbally categorize learned items as familiar or novel^1–3^. However, relying exclusively on verbal reports entails significant limitations. First, from a basic science standpoint, it confounds memory per se with the ability to access and report the memory. Accordingly, it remains unclear whether the underlying neural process, a deleterious effect of brain disease, or the benefit of sleep pertain to memory, or to the ability to access or report the memory engram. In addition, studying memory retrieval is limited (or completely impossible) in populations where explicit reports are unreliable or absent, such as in aphasia patients, newborns, or animals. A reliable and well-validated method to assess episodic-like memory of events without an explicit report would represent a major advance. By “episodic-like memory”, we refer to computational definitions (e.g. ^4–6^) that emphasize a memory for an episode (an event of what, where, and when) irrespective of the autonoetic (personal) and conscious aspects of the memory (as in the full definition by Tulving ^3^).

A similar challenge exists in the study of consciousness, where specialized paradigms have been developed to separate the neural correlates of consciousness from those related to its report^8^. Many such paradigms employ eye tracking, suggesting that an eye-tracking-based “no-report” paradigm could also be effective in studying memory retrieval. Additional motivation arises from rodent studies, where differences in exploratory behavior are routinely used to index memory^8,9^. Humans and other primates are visual-centric and primarily rely on eye movements to explore their environment^10–12^. Along this line, Kano & Hirata (2015)^13^ leveraged visual exploration to assess memory retrieval by tracking great apes’ eyes during a single movie clip. Inspired by this approach, we sought to use the exploration profiles of human gaze in a similar way, which could potentially allow to capture individual memory traces independent of explicit reports.

Already over the past two decades, eye tracking has been increasingly linked to human long-term memory^14^. To this end, a variety of studies in the fields of in visual search^15,16^ and spatial orienting of attention^17,18^ have examined memory-guided eye-movements. Contextual cueing paradigms^19^, visual expectation paradigms^20^, and “looking-at-nothing” paradigms^21,22^ have jointly shed light on how declarative^23^ and non-declarative^18,24^ memory can guide human gaze. It is therefore widely accepted that people visually explore static images in relation to the retrieval of memories^25–32^. For example, the recognition of photographs is associated with a decrease in distributed overt attention, and familiar faces elicit fewer fixations compared to unfamiliar ones^29,30,32–34^.

We aimed to leverage gaze memory effects to develop a paradigm and analytical approach that uses eye-tracking as an alternative to verbal reporting. To this end, we developed and validated a method that uses *anticipatory gaze* patterns to quantify memory of events. We hypothesized that predictive eye movements during video clip viewing could serve as a proxy for the retrieval of episodic-like memory in adult humans. Due to its non-verbal nature, anticipatory gaze has previously been used to study memory in preverbal infants^35–37^ and toddlers^38^. Recently, memory-guided predictions, reflected in anticipatory gaze behavior, have regained attention in memory research^39–43^. Such studies connect predictive eye movement patterns to memory updating^42^, to scenes with potential to change^40^, or to tracking the transition from learning to memory-guided action^39^. However, it remains unclear how reliable anticipatory gaze can be used as an indicator of episodic-like memory in adults and how such effects relate to explicit reports of episodic-like memories.

Here we introduce and validate MEGA (Memory Episode Gaze Anticipation), a no-report method based on eye tracking during repeated movie viewing to assess memory for events. Participants watched movies with predefined surprising events. When participants watch the movies for the second time and have already formed a memory of the event, their gaze is drawn to its location, anticipating its occurrence before it appears. Therefore, the movies deliberatively produce an anticipatory gaze, similar to a cued recall task. Importantly, this design allows the movies to be presented twice in exactly the same way, with memory being the only difference between the two viewings. In a series of experiments, we validate MEGA as an approach to quantify memory without report and compared it to explicit memory retrieval. We introduce a distance-based metric on eye-tracking data to capture gaze characteristics probing memory at the single-trial level. We then compare its correspondence with multiple explicit memory reports and contrast it with other eye-tracking metrics, such as pupillometry. Finally, we demonstrate one application by studying the influence of sleep on memory consolidation via MEGA without explicit reports.

## Methods

### Participants

We tested a total of 145 participants across the five different experiments. Written informed consent was obtained from each participant prior to their involvement in the study as approved by the Institutional Review Board at Tel Aviv University (Experiments 1,2) or by the Medical Institutional Review Board at the Tel Aviv Sourasky Medical Center (Experiments 3,4). All participants were required to have normal or corrected-to-normal vision, reported overall good health, and confirmed the absence of any history of neurological or psychiatric disorders. Experiment 1 (animation movies) included a total of 34 participants (age range: 19-61, M ± SD = 26.2 ± 9.01 years; 23 (67%) female participants and 11 (33%) male participants). Experiment 2 (animation movies - with elaborate memory assessments) included 32 participants (age range: 18-40, M ± SD = 26.48 ± 4.3 years; 23 (72%) female participants and 9 (28%) male participants) of which we excluded two due to unsufficient eye-tracking. Experiment 3 (naturalistic movies) included 32 participants (age range: 19-44, M ± SD = 27.03 ± 5.09 years; 9 (27%) female participants and 23 (73%) male participants), where one participants eye-tracking failed and was subsequently excluded. For experiment 4 (naturalistic movies - with sleep consolidation), 19 of 27 participants reached sufficient sleep (sleep efficiency>50%). These were subsequently analyzed (age range: 20-34, M ± SD = 27.13 ± 3.44 years; 10 (34%) female participants and 9 (66%) male participants). For the control experiment animation – no SE), 20 participants were subsequently analyzed (age range: 23-51, M ± SD = 28.8 ± 6.88 years; 14 (70%) female participants and 6 (30%) male participants). Thus we collected 145 data sets and analzed the data of 134 participants after the exclusion of 11 participants. Information about sex was provided by participants’ self-report, we did not collect gender.

### Experimental Procedures

Five different experiments were carried out, each included a first viewing session (encoding), a break (consolidation period with different durations), and a second viewing session (retrieval) of the same movies, and in some cases additional new movies.

**Experiment 1: Tailor-made animation movies.** Experiment 1 (Figures 1, 2, 3) was conducted during a daytime session with a two-hour break between the first and second movie viewing sessions. Following the completion of consent forms, and eye tracking setup (see below), the first viewing session started around noon (11:48 AM on average) and lasted an average 45 minutes. Participants watched 65 animated movies while pupils and gaze were monitored (see below). The movies were separated by a 2-second fixation cross presented on a blank grey screen. Then, the participants had a ∼ two h break in which they were free to leave the lab unsupervised. During the second viewing session, each movie was followed by participants feedback indicating their explicit memory recall (“*Have you seen this movie before?*” with options Yes (1) or No (2)) and a second screen in which they rated their confidence (“*How confident are you in your answer?*” on a scale from “*Not at all (1)*” to “*Very confident (4)*”).

**Figure 1.**
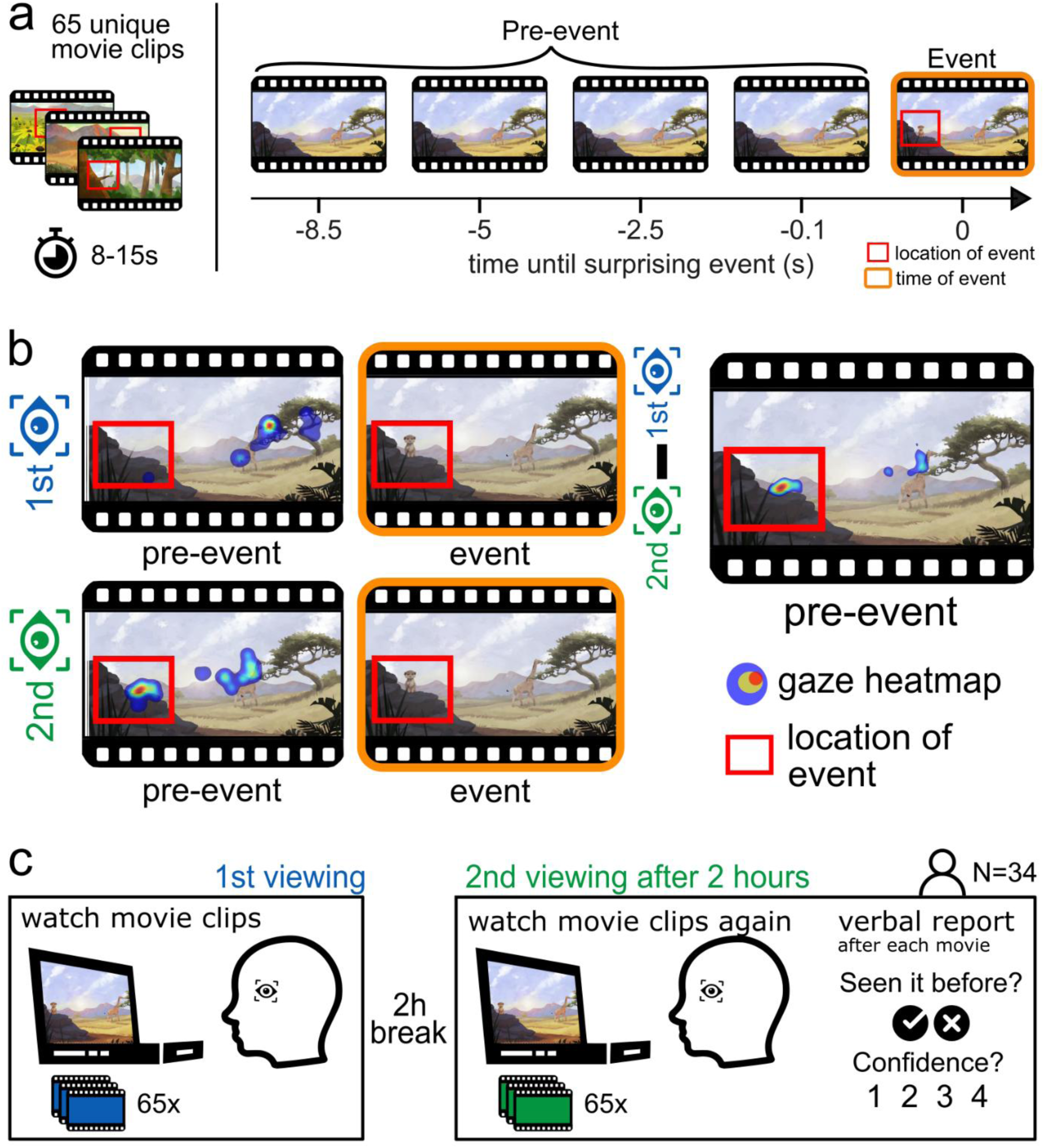
MEGA paradigm. (a) Visual Stimuli. 34 Participants watched 65 custom-made movie clips, each featuring a distinct surprising event. Each surprising event included a specific object at a specific location (red square) at a specific time (orange square). (b) Analysis rationale: gaze patterns were compared during the 1st and 2nd viewings to uncover memory retrieval. Top row: the average gaze location during 1^st^ viewing (blue); Middle row: the same average gaze during the 2^nd^ viewing (green). Bottom row: the difference between the gaze during 1^st^ and 2^nd^ viewings before the event appears on the screen. Participants were expected to gaze more often toward the location of the upcoming event, indicating their memory of the event. Heat maps represent gaze locations of a single move, averaged over time. (c) Experimental design: 65 movie clips were presented twice in randomized order, with a consolidation interval of 2 hours between viewings. Eye-tracking data was collected during both viewings. During the 2^nd^ viewing, after each movie, participants reported whether they recognized the movie clip from the first session and rated the confidence of their response.

**Figure 2.**
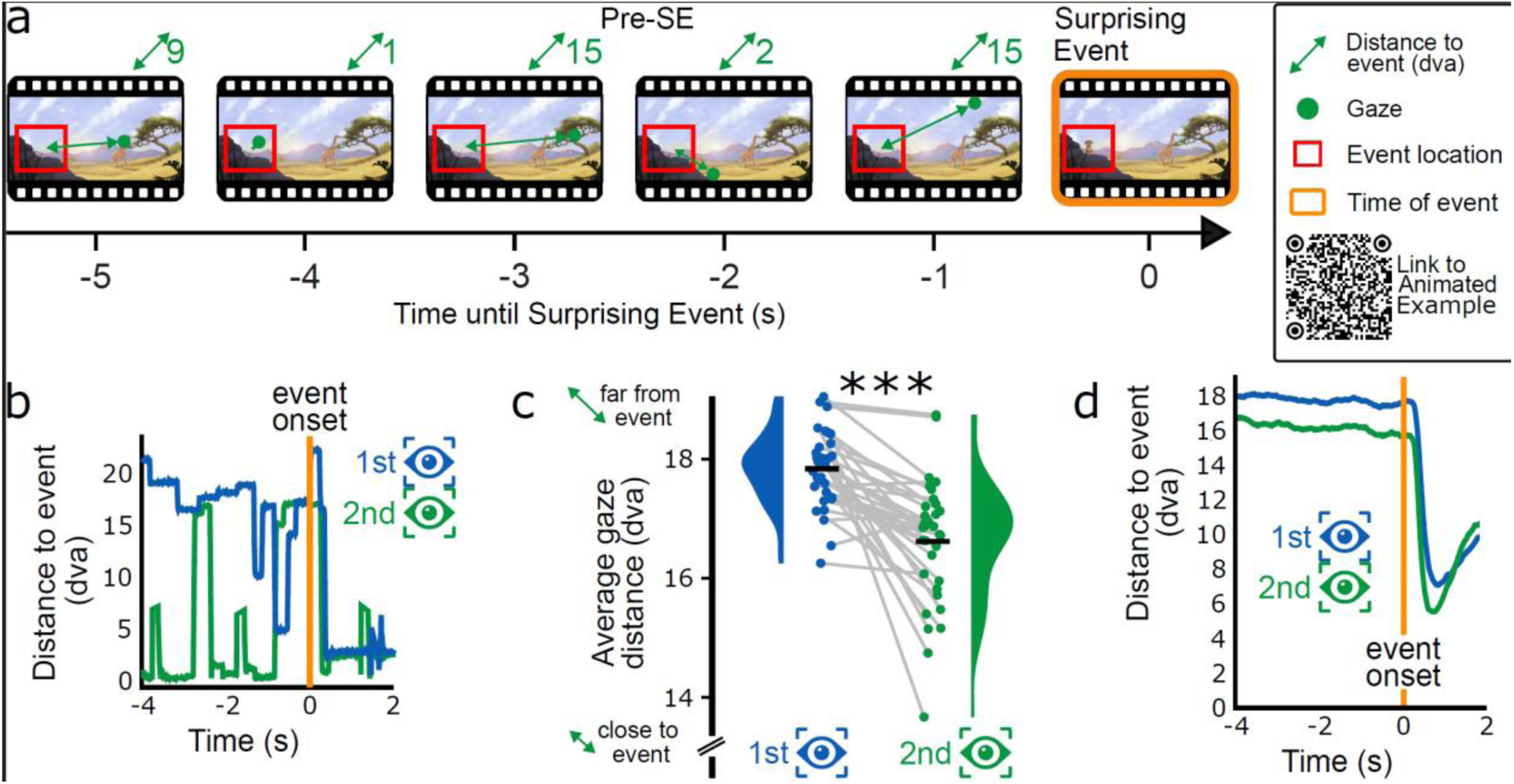
Anticipatory gaze effect. A) Gaze Proximity Measurement: participant’s gaze trajectory while watching a movie clip. The Euclidean distance to the location of the surprising event center point is calculated at each time point in Degrees of Visual Angle (DVA). The QR code leads to an animated illustration of the methodology (for example, a movie with superimposed eye movements and gaze distance measures). B) Temporal Gaze Distance of a single participant during first (blue) and second (green) viewings of a clip. The SE’s onset is marked by an orange line, serving as a temporal marker for gaze behavior. C) Gaze distance (GAD) comparison within participants of Experiment 1 across two viewings (N=34). The gaze average distance is calculated relative to the event center prior to its appearance on the screen and averaged across movies. GAD significantly decreases during the second viewing compared to the first viewing indicating memory for the event’s location. Black horizontal line: group average, colored area: density estimate of GAD distribution, dots: individual participant average, line connects 1st and 2nd viewing of the same participant. D) Averaged gaze temporal dynamics: An aggregate view of gaze distance from the event across all participants and movies, time-locked to the event onset. The distances as a function of time for the first (blue) and second (green) viewings show the participants’ anticipation of the event based on their prior viewing experience. Participants were surprised and looked at the surprising event immediately after its appearance, which can be seen in the steep slope after the event onset (orange line). *** = p<0.0001

**Figure 3.**
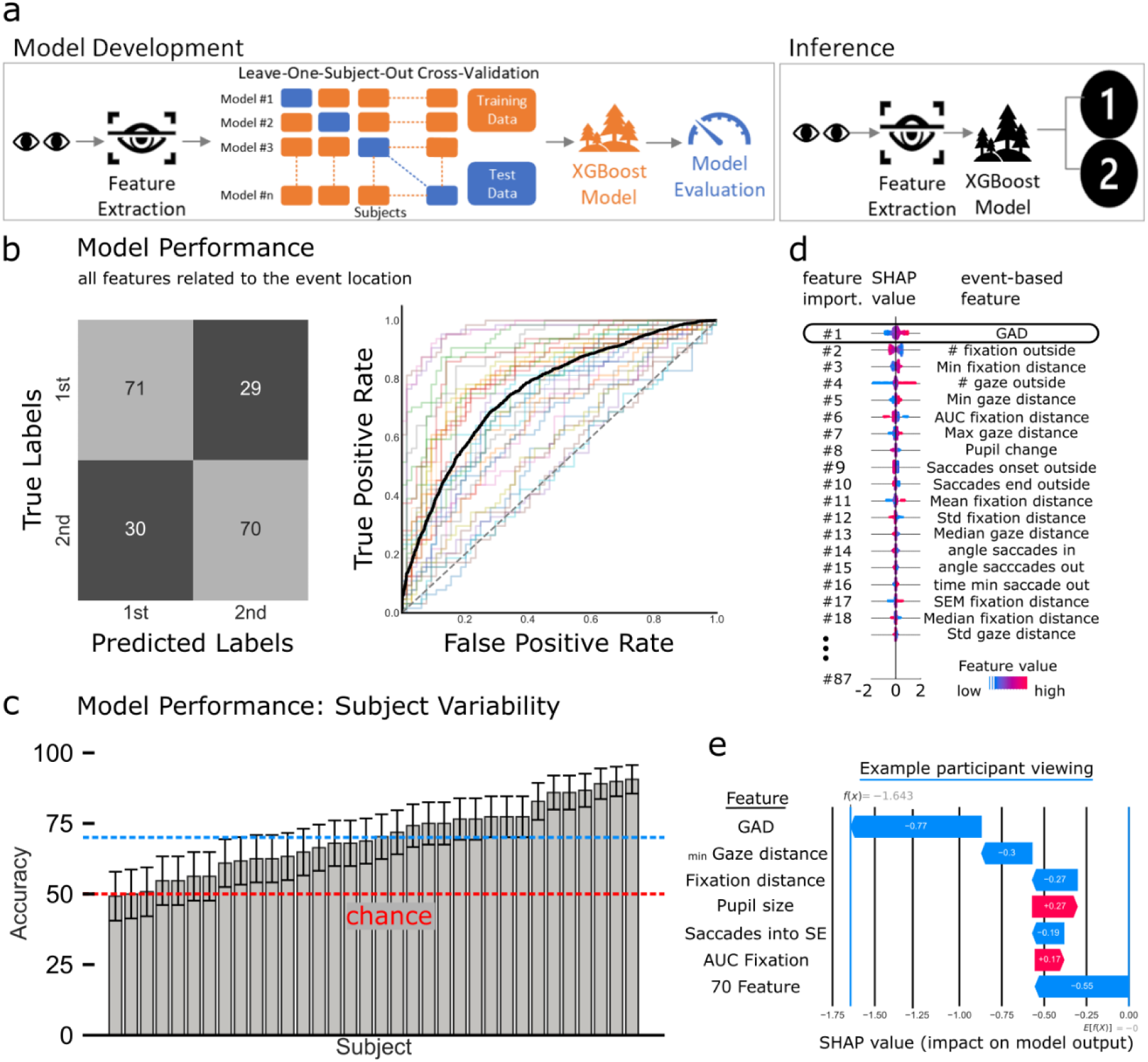
Single-trial Predictive Modeling Using Machine Learning of Eye Tracking Data Features. (a) Model Development and inference: We utilized an XGBoost classifier, leveraging 87 features derived from the event location and timing to perform single-trial level classification. The model’s robustness was ensured through a leave-one-subject-out cross-validation technique, where the algorithm was iteratively trained on the dataset excluding one subject, which was then used for testing. (b) Confusion Matrix: The average confusion matrix across all subjects (N=34), summarizing the classification performance of XGBoost models. Values along the diagonal (top left and bottom right) entries (71%,70%) represent the percentage of correct classifications, indicating true positives and true negatives, respectively. Off-diagonal values (bottom left to top right) entries (30%, 29%) denote misclassification probability. The Receiver Operating Characteristic (ROC, on the right) curves for XGBoost models, each tested on a distinct subject within our study cohort. These curves plot the true positive rate (TPR) against the false positive rate (FPR) at various decision threshold levels. Each trace represents the ROC curve for a subject-specific model (different color for each participant), delineating the trade-off between sensitivity and specificity. The black line denotes the mean ROC curve across all models. (c) Performance across individual participants (bottom left): bar chart displaying the classification accuracy for each subject, with the average accuracy across all subjects (blue, 70%) and chance-level (red, 50%). Error bars signify confidence intervals reflecting the precision of the model for each subject. (d) SHAP feature analysis: ranking of the influence of various gaze metrics on the model’s predictions. Out of 87 SE-based features, the distance-based features, particularly GAD, emerged as the most important feature. (e) A waterfall plot provides an example of how individual features contribute to a single trial’s classification.

**Experiment 2: Animation movies with extensive explicit memory reports.** This experiment (Figure 4) examined in more detail the relation between anticipatory gaze and verbal reports, probing different aspects of memory. The experiment employed the exact same animation used in Experiment 1 for the first viewing. However, in the second viewing, the participants viewed an edited version of these movies where the surprising event E was omitted from each movie. In the first viewing, participants watched 48 movies. After a 2h break, participants were instructed about the tasks associated with the second viewing ahead and a single training example verified they could successfully follow the tasks. During the second session, participants watched the edited, event-lacking version of the 48 movies and 12 new movies in a random order. After each movie, the following five retrieval tasks were presented:

1. **movie recognition:** “Do you remember watching this movie before?” (“Yes” or “No”),
2. **free recall:** “Please describe what was missing in the video and where did it happen?” Participants were instructed to say their answer aloud (for example: “snake on the right” or “frog in the center”).
3. **object recognition:** two objects were presented, and the participant was asked to indicate, “What was missing in the movie?”. The lure object was created such that it was not unlikely to exist in the presented environment. For example, if the scene was underwater, possible lures would be “fish” or “crab” but not “elephant”. Hence the lure list was fixed in such a way that each object appeared once as correct and once as incorrect answer.
4. **event location recall:** a frame from the movie was presented, and participants were instructed to click on the screen where they thought the event should have happened. An answer was considered correct if they clicked in the correct quarter of the screen.
5. **temporal recall:** participants were asked about the timing of the event within the Movie: “When did the Surprising Event happen?”. Answer options were a) “In the middle of the movie” or b) “In the end of the movie”. Event timing was considered in the middle or in the end according to the distribution of all events. If an event appeared before the median time of all SE’s it was considered in the middle, and vice versa.

**Figure 4:**
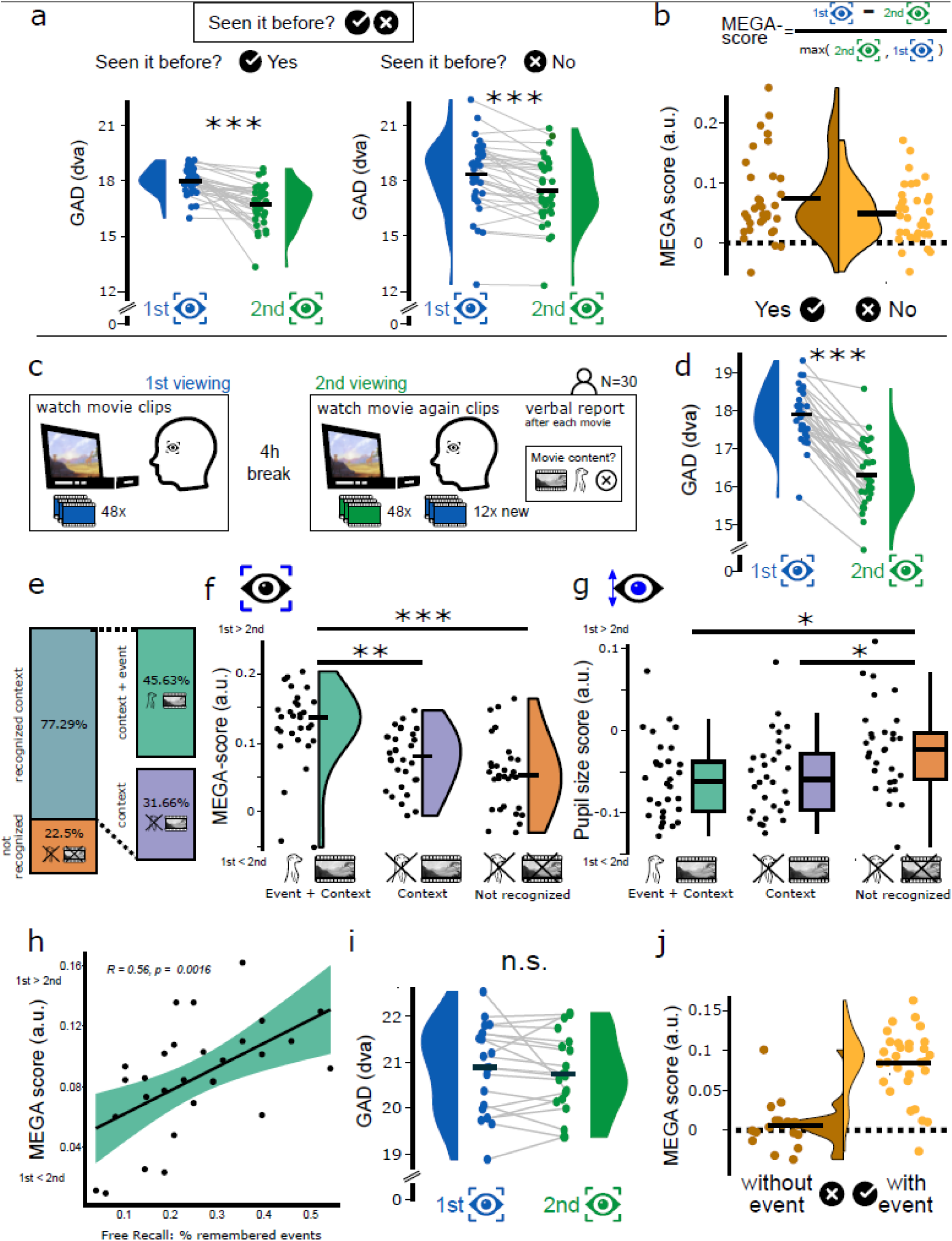
The relationship between anticipatory gaze and explicit memory reports. A) Experiment 1 (N=34): gaze in the second viewing of movies explicitly reported as “seen” (green) showed greater proximity to the event location compared with the first viewing (blue) for both recognized and unrecognized movies, signifying memory retrieval. B) The MEGA score quantifies the change in GAD between the first and second viewings, calculated using the formula above. This MEGA score is higher for movies that were explicitly remembered versus not remembered. C) Procedure of the Experiment 2 (N=30): Participants completed a similar task except surprising events were omitted from the second viewing and followed by an extensive explicit memory report. Memory was assessed to determine what exactly participants remember. D) Replication of the anticipatory gaze effect: consistent with experiment 1, anticipatory gaze appears in the second viewing (green) and not in the first viewing (blue) across all naïve 30 participants. E) Categorization of explicit report. Participants did (i) not recognize the movie clip at all (orange), (ii) recognize the context alone (violet), or (iii) recognize the context and recollected the event (green). F) The MEGA score is higher for movies in which participants remembered the event in full, compared to those with only scenery memory or completely forgotten movies. There is no significant difference between movies with scenery recognition and movies that were not recognized. G) Pupil size variation and memory: Examines the decrease in pre-event pupil size from first to second viewing, in relation to scene recognition and event recollection, highlighting smaller changes (1^st^ vs 2^nd^ viewing) in unrecognized cases. H) The MEGA score correlates with participants’ event recollection in the free recall. Every dot represents a participant. I) If we remove the events completely from the experiment, anticipation diminishes and gaze in the second viewing (green) shows comparable proximity to the event location compared with the first viewing (blue). (J) MEGA score for the control experiment without any events, in comparison to MEGA score of Experiment 1 (same procedure for both experiments), revealing chance-level MEGA scores for experiments with movies without events. Plots represent the median (bold horizontal line), 95% confidence intervals (whiskers), density plots, and subject averages (dot plots). * = p<0.05, ** = p<0,01, *** = p<0,001.

The free recall was self-paced, whereas the other four questions were limited to 5 seconds. The movie order in the first and second viewing was randomized, but which movies were presented once or twice was fixed and identical for all participants.

**Experiment 3: Naturalistic movies.** Experiment 3 (Figure 5) investigated anticipatory gaze at our sleep lab using naturalistic videos from YouTube. After setting up the eye tracking and EEG (below), the first viewing session started around 2 PM and included watching 100 naturalistic movies. Then, the participants had a 2-hour break and were instructed to remain awake. The second viewing session also included 100 movies (80 seen movies and 20 new ones) as well as a simple recognition task (“*Have you seen this movie before?* (“Yes” or “No”),”) and confidence feedback (“*How confident are you in your answer?”* on a scale from “*Not at all (1)*” to “*Very confident (4)*”)”) as in Experiment 1. Since the movies were publicly available on YouTube, there was a possibility that the participants had already seen some of them before the experiment. To address this, a recognition task was also carried out during the first viewing, and any trials with positive responses were excluded.

**Figure 5:**
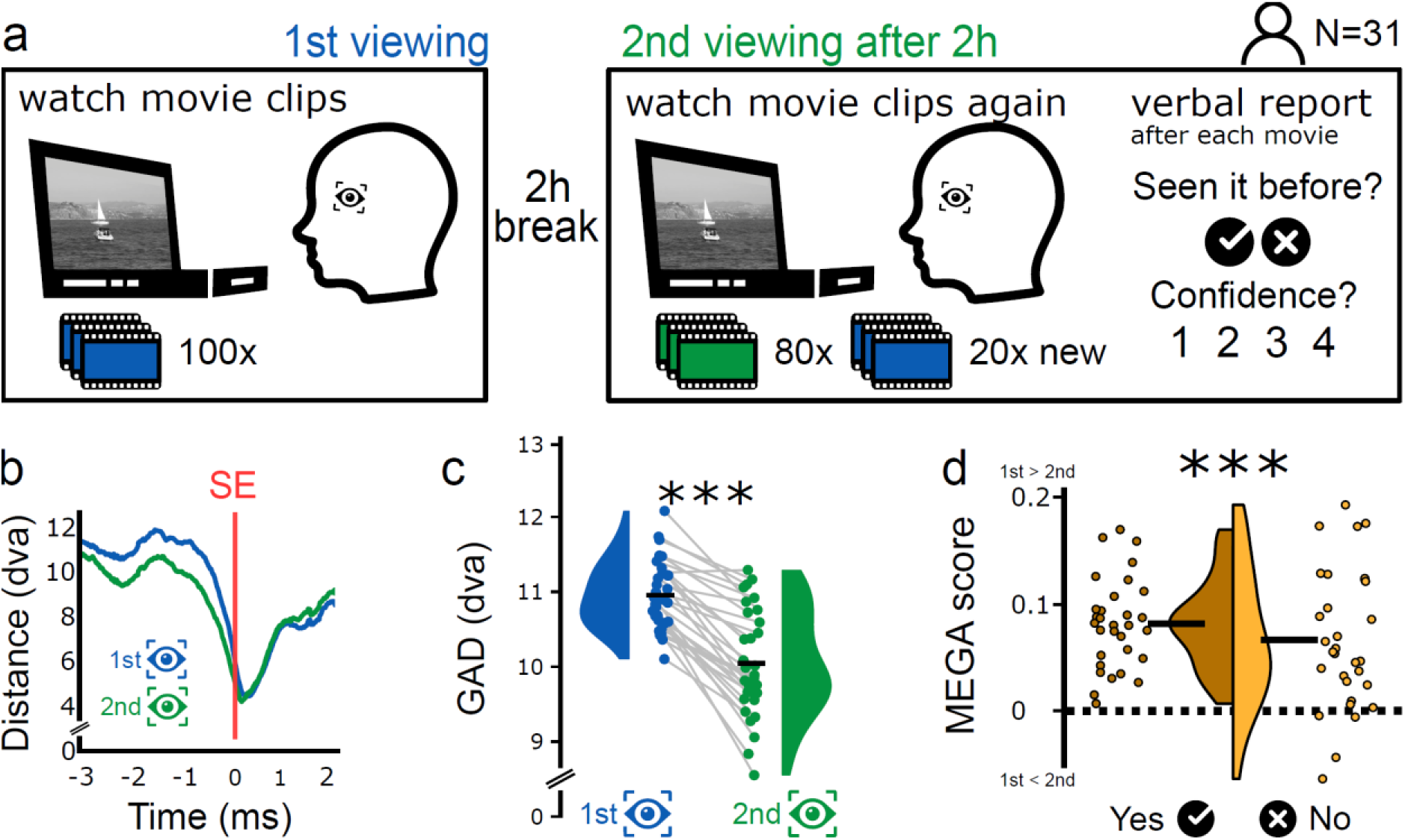
The anticipatory gaze effect is replicated in naturalistic (YouTube) videos. (a) Experimental procedure: 32 participants viewed 100 YouTube movies in two sessions spaced 2 hours apart. In the 2^nd^ session, 20 novel movies were included along with 80 of the original movies, and verbal reports were collected after each movie. (b) Average gaze distance to the surprising event for 1^st^ (blue) and 2^nd^ (green) viewing. A drop in gaze distance that already begins before event timing shows some anticipation of the surprising event in naturalistic movies, probably due to narrative and camera movements. (c) GAD decreases in 2^nd^ viewing (green) compared to 1^st^ viewing (blue): p = 9.3e-10, Wilcoxon Signed-Rank Test). (d) MEGA scores as a function of explicit memory reports, showing a higher MEGA score for movies that participants remember (p<0.0001).

**Experiment 4: Naturalistic movies with nap or wake.** In Experiment 4, the setup was identical to Experiment 3 but introduced new participants and a modification: participants were given a nap opportunity during the 2h break while we monitored EEG, electrooculogram (EOG) and electromyogram (EMG) to monitor sleep. Data collection and sleep scoring were previously described^44^. Data from 8/27 participants was excluded due to short sleep duration (<50% sleep efficiency).

**Control experiment: Animations without an event.** In a control experiment we repeated experiment 1 but the surprising event was removed in both the first and second viewings. Therefore, any change in gaze between the viewings would reflect changes based on familiarity with the scenery.

**Figure 6.**
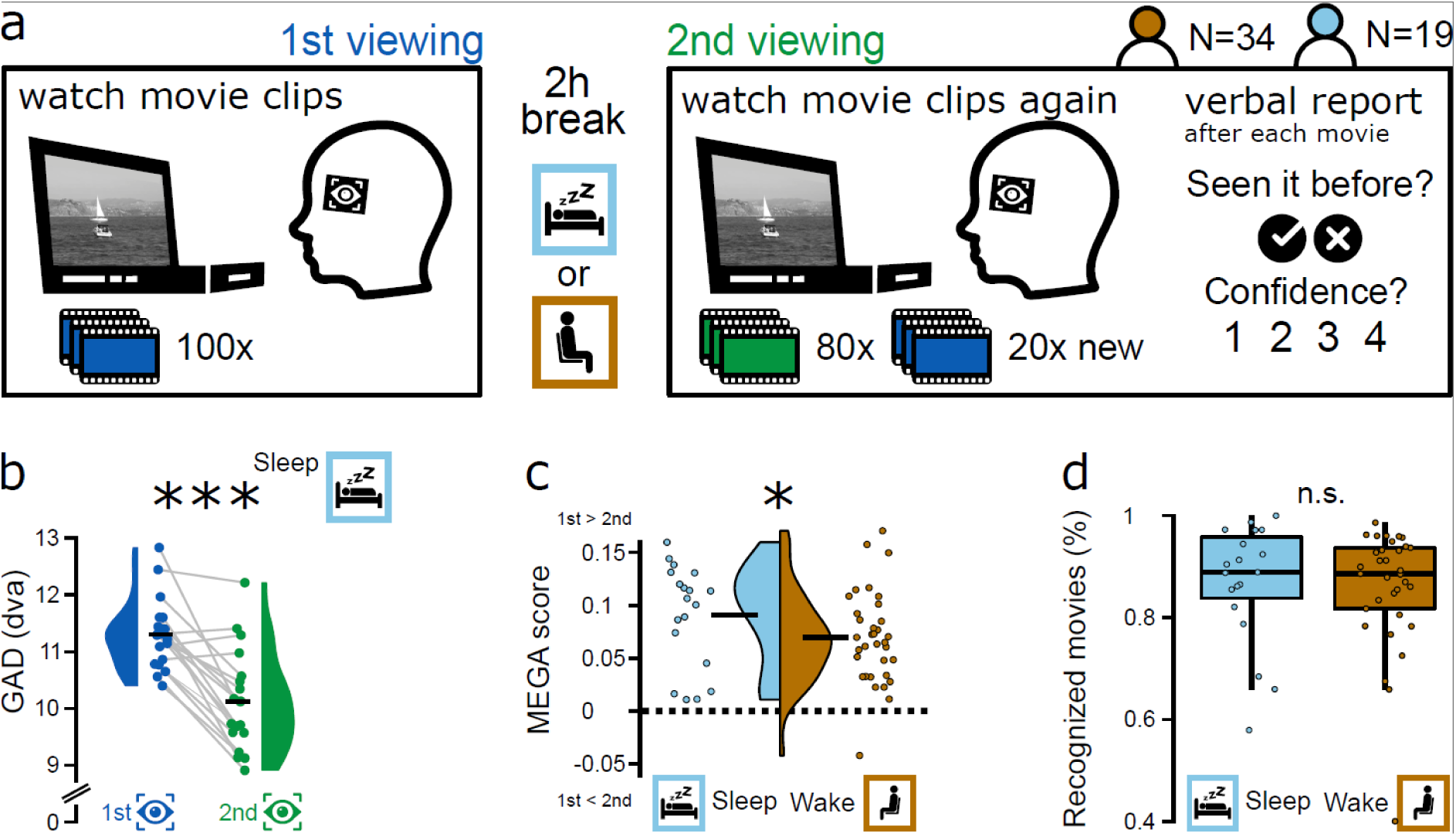
MEGA scores improve after sleep compared to wakeful rest. (a) Experimental design: participants (N = 19 for nap condition; N=34 for wakeful rest condition) watched naturalistic videos as in a “naturalistic” experiment with YouTube videos. The 2-hour interval between the 1^st^ and 2^nd^ viewing included either a nap opportunity or wakeful rest. (b) GAD in 1^st^ (blue) and 2^nd^ (green) viewing for the nap condition participants reveals a significant decrease in GAD during 2^nd^ viewing (p = 9.3e-10, Wilcoxon Signed-Rank Test) (c) A comparison of MEGA scores (normalized difference in GAD between 1^st^ and 2^nd^ viewings) in the wake (brown) and sleep (cyan) conditions reveals greater MEGA after a nap (p = 0.0439, Wilcoxon Signed-Rank Test). (d) Comparison of the recognition rate in the wake (brown) and sleep (cyan) conditions do not show significant differences in the verbal report (BF_10_ = 0.390± 0%). Box plots represent the median (bold horizontal line), the interquartile range (box, 50% of data) and 95%confidence interval (whiskers).

### Movie stimuli and visual presentation

Overall, we used 185 different movie clips. We presented movies in full-screen mode to maximize engagement. Between the movie clips, a gray background with a fixation cross was displayed for 2 seconds to standardize the visual field and prepare the participants for the upcoming stimulus. In all experiments the order of the movie clips was pseudo-randomized for each participant to avoid order effects that could influence memory encoding and retrieval processes. Previously unseen movies were incorporated in the second session to increase task difficulty and enhance performance variability but were not subsequently analyzed. The experiments were coded in Python, using the PsychoPy^45^ package and the PyLink package, which facilitates interfacing with EyeLink eye-trackers (see below).

For Experiment 1 and 2, we used 65 silent animated colored movie clips. For the naturalistic movies, we included 100 silent black and white movie clips collected from YouTube. Detailed description and examples can be found in Supplementary Information (Supplementary Movie S2 and Supplementary Figure S3).

### Eye tracking

Eye tracking employed EyeLink 1000 Plus (SR Research) as in Sharon et al.^46^ with a sampling rate of 500Hz. We first determined the dominant eye of each participant, utilizing a modified version of the Porta test^47,48^. Participants were then instructed to position their heads on a chin rest 50-70cm from the screen to maximize eye tracking quality. Next, a 9-point calibration and validation process was performed until the error was below 0.5° of visual angle.

**Event-Related Analysis.** The gaze coordinates and pupil size were computed for each movie seen by the participants. A custom validation tool was employed to ensure data integrity, running several checks such as for the correct number of files and recording rates (code available). Movie trials with more than 30% tracking loss were excluded. Furthermore, all movies that were not shown in the first encoding session were excluded as well (12 animations & 20 naturalistic videos). Next, we computed the Euclidean gaze distance between each gaze coordinates and the center point of the event location (in degrees of visual angle, DVA). Data quality was visually ensured, example can be found in the Supplementary Movie 1 & 2.

**Gaze Average Distance (GAD).** GAD quantifies the mean Euclidean distance between each gaze data point and the SE center point, calculated separately for each time point prior to the appearance of the SE and averaged across all computed distances (not just fixations). Thus, GAD captures a cumulative estimate of how closely participants’ gazes approximate the surprising. GAD was averaged for each subject across all movies within each session.

**MEGA Score: a metric for gaze anticipation.** The MEGA Score is a normalized metric reflecting how anticipatory gaze behaviors change across repeated viewings of the same movie: MEGA Score = (GAD^1st^ ^viewing^ - GAD^2nd^ ^viewing^)\ max(GAD^1st^ ^viewing^, GAD^2nd^ ^viewing^). Because the averaging of ratios is biased, specifically if the denominator changes, we divided by either the average of the 1^st^ viewing or the average of the 2^nd^ viewing, depending on which was larger. Importantly, central outcomes remain consistent regardless of whether normalization was applied, or which specific normalization method was used (no normalization, normalization by the max, by the baseline and by the mean). Higher MEGA Scores indicate that participants looked closer to upcoming SEs during the second viewing, reflecting enhanced anticipatory gaze behavior guided by memory.

**Pupillometry.** Pupil size was extracted during all fixations prior to the surprising event using the EyeLink segmentation tool. Trials with >30% missing data were excluded. Pupil size prior to the events was used to compute a normalized score (as for the MEGA score): pupil size score = (pupil size ^1st^ ^viewing^ - pupil size ^2nd^ ^viewing^) / max(pupil size ^1st^ ^viewing^, pupil size ^2nd^ ^viewing^). The pupil size score provides a nuanced measure of change in pupil size across repeated movie viewings, where a higher pupil size score reflects a larger pupil size in the 2^nd^ viewing.

### Behavioral analysis

To disentangle the relationship between anticipatory gaze and explicit report, we collected verbal reports in all four experiments. For Experiment 1, 3 and 4 we asked after each movie “*Have you seen this movie before?* (“Yes” or “No”),” and collected their confidence rating. In Experiment 2, five retrieval tasks were presented: i - movie recognition, ii - free recall, iii - object recognition, iv - location recall, v - temporal recall. For the anticipatory gaze analysis in relation to the explicit report, we computed the MEGA score for each movie. To estimate the difference between the first and the second viewing, we averaged the MEGA score over all movies for each participant (‘new’ movies were excluded) and then tested the MEGA score against chance. To estimate the subsequent memory effect, we compared the average recognized and unrecognized MEGA score. This was done for naturalistic and animated movies. Moreover, we used the same procedure to compare the sleep and the wake groups. In Experiment 2, anticipatory gaze was evaluated in relation to event recollection. Based on the four forced-choice tasks (i – movie recognition, iii – object recognition, iv – location recognition, v – early or late SE), we defined and evaluated three labels of remembered memory content: (A) context and event recollection, (B) context recognition, and (C) not recognized. The free recall (ii) was analyzed separately. An extensive description of the process and the result of each task is available in as Supplementary Figure S1.

### Single-Trial Decoding of Movie Viewing

Raw eye-tracking data were transformed into 243 engineered features containing fixation, saccades, blinks, and pupil related parameter. 87 of these parameters were in relation to the location and timing the surprising event (see Supp. S7 & S8). An ensemble of XGBoost classifiers was employed, optimized using grid search, and evaluated with leave-one-subject-out (LOSO) cross-validation to classify first and second viewings of the movies. Model performance was assessed using classification accuracy, confusion matrices, and ROC curves. The statistical significance of the classification was quantified against the chance level (50%) by computing a one-sample t-test of the ROC AUC score. SHAP analysis was applied to interpret the contributions of individual features to predictions, providing insights into the model’s decision-making process.

### Statistical analysis

None of the reported experiments were preregistered. We used parametric methods for statistical testing of normal data. Specifically, we computed the suitable t-tests and Cohen’s D for direct comparisons. For the comparisons of three groups (event memory, scenery memory, no memory) one-way ANOVAs were computed and Effect sizes were estimated using eta-squared (η²), which represents the proportion of variance explained by the independent variable. Post hoc comparisons were conducted using Tukey’s Honestly Significant Difference (HSD) test following a one-way ANOVA to identify pairwise differences between group means. In Experiment 2 we corrected for multiple comparisons when examing all teh verbal report tasks. Where applicable, normal distribution and equal variances were formally tested. For non-normal data, we used Wilcoxon signed-rank test and rank-biserial effect sizes from Wilcoxon-rank test. The statistical significance of ML classification was quantified by applying a one-sample t-test to the ROC AUC score compared to the chance level (50%). To evaluate the evidence for or against the null hypothesis, we conducted Bayesian null hypothesis testing by computing Bayes factors (BF₁₀), quantifying the relative likelihood of the data under the alternative versus the null model. The prior distributions used to compute BF₁₀ were informed by effect sizes from the most methodologically similar experiment. Pearson’s correlation coefficient was used to assess the linear relationship between variables. All t-tests were two-sided, except for the sleep–wake comparison, which was one-sided due to a strong a priori expectation that sleep would enhance memory retrieval. The error probability of 5% was chosen for all statistical tests and the statistical tests were either computed in R or Python. The sample size of Experiment 1 was similar to those generally in the field. All the following experiments were estimated with a power analysis based on the effect size of experiment 1 (Cohen’s D = 1.8).

## Results

We constructed the Memory Episode Gaze Anticipation (MEGA) paradigm with the aim of capturing memory retrieval in its raw form without the additional layer of verbal reports. Participants viewed short (8-23s) movie clips in two sessions conducted a few hours apart while gaze was monitored using an infrared video-based eye tracking system (see Methods). Each movie contained a surprising event that saliently occurred in an unexpected location and time (e.g., an animal suddenly appearing behind a rock, Figure 1A; see A1 for the full movie). The movie clips were custom-designed animations depicting simple ecological scenarios. Precise event timings and locations were equally distributed across the movie length and screen space (see methods). We hypothesized that in the first viewing, gaze patterns before the event occurs (‘pre-event) would be exploratory, whereas, in the second viewing, after the formation of memory, gaze patterns would anticipate the event and preferentially occur around the event location (Figure 1B). Accordingly, the memory for the surprising event may manifest as a difference between the 1^st^ and 2^nd^ viewings at pre-event intervals around the anticipated location of the event (red rectangle and heat maps in Figure 1B). To test this, the first experiment included 34 participants who viewed 65 movie clips in two viewing sessions separated by a two-hour break (Figure 1C and see detailed methods). In the 2^nd^ viewing session, verbal reports and confidence measures were collected after each movie, to allow comparison with standard explicit reports.

### Gaze anticipation indexes memory for events

To quantify anticipatory gaze towards the remembered location in each movie viewing, we assessed, for each time point separately, the Euclidean distance from the gaze location to the event location (a priori defined by the animation studio that compiled the movies, Methods). Figure 2A illustrates this calculation, which utilized all data points (not only fixations) to transform the multivariate eye tracking data into a single time series that encapsulates anticipatory gaze behavior. During the 1^st^ viewing, the distance from the event location was mostly large, but upon 2nd viewing, we observed that the gaze gravitated toward the expected location of the event before its onset (see representative example of a single movie in Figure 2B). We quantified this by the Gaze Distance (GAD), the mean distance from each gaze point to the event location from the movie’s beginning until the event onset. GAD is a simple indicator that can reveal a tendency to gaze closer to the event location before its onset. Computing the GAD across all movies for each participant (Figure 2C) revealed a significant convergence to the event location upon 2^nd^ viewing, observed in 31/34 (91%) of participants (mean GAD 1^st^ viewing=18.13±0.62° vs. mean GAD 2^nd^ viewing=16.91±1.05°; t(33)=6.199, p= 5.377e^−07^; Cohen’s d = 1.4,, 95% CI [0.87 1.84]). To test if implicit learning may play a role, we compared the GAD for the first ten movies to the last movie in the 1^st^ viewing session. GAD values were comparable (first ten vs last movie: BF_10_ = 0.20 ± 0.06%, prior according to subsequent memory effect, see below), suggesting that effects were not driven by implicit learning of the surprising event’s appearance. Next, to observe the temporal dynamics of anticipatory gaze, GAD was averaged across all movies and participants without averaging across time (Figure 2D). This analysis revealed a closer gaze towards the event location in the 2^nd^ viewing that was present throughout the seconds leading to its appearance (Figure 2D, green time-course), followed by a sudden drop in GAD after-event appearance (reflecting gaze towards the surprising event once it occurred). The anticipatory effect, measured as the difference in GAD between the first and second viewing, remained stable over the entire pre-event interval, without evidence for a temporal reinstatement specifically timed to the event (Supplementary Figure S4).

### Machine learning discriminates first and second viewings at a single trial resolution

We tested to what extent machine learning (ML)-based classification of multiple eye tracking features could extend the intuitive GAD metric and accurately identify memory traces at the single-trial level. First, we focused on reducing gaze distance as captured by the GAD metric (above). We employed the XGBoost classification algorithm^49^ at the single-trial level, complemented by a leave-one-subject-out cross-validation method (Figure 3A). Models achieved an average correct classification of 69±10% (chance: 50%), demonstrating the model’s ability to distinguish, based only on GAD, whether a single trial’s data represented the first movie viewing or whether it had been viewed before-1^st^ or 2^nd^ viewing (Figure 3B). The Receiver Operating Characteristic (ROC) curves for each subject-specific model and the average ROC curve indicated consistent model performance across subjects (Figure 3C). Accordingly, the area under the ROC curve (AUC-ROC) was 0.75±0.26, quantifying the model’s overall effectiveness (one-sample t-test: t(33) = 10.76, p = 2.47e-12). In 32/34 of the participants (94%), model accuracy was greater than chance levels, with an average accuracy of 0.69±0.1, attesting to its reliable prediction of identifying memory traces (Figure 3D).

Next, we employed an exploratory ‘bottom-up’ ML strategy by broadening our analytical scope to encompass a wider array of eye-tracking features, irrespective of our initial GAD metric. We manually engineered multiple features from eye-tracking data centered around the event location, thereby minimizing potential session-specific confounds such as differences in time-of-day and cognitive load due to tasks and reports in 2^nd^ viewing (Methods). Such features (87 in total) included aspects such as fixation count ratios relative to the event location, the velocity and visual angle of saccades directed towards the event, and the change in pupil radius during fixations within the event location compared to prior fixations, thereby capturing event-related eye tracking spatial and temporal features. With these features, models achieved an average classification accuracy of 71±11.73% and 70±12.34% for 1^st^ and 2^nd^ viewings, respectively. Associated AUC-ROC metrics exhibited an average of 0.75±0.26 (one-sample t-test: t(33) = 10.85, p = 1.98e^−12^). In 32/34 of the participants (94%), model accuracy was greater than chance levels, with an average accuracy of 0.7±0.11. Remarkably, our initial analyses based solely on the GAD metric already showed comparable performance to the extensive 87-feature model (see Supplementary Notes 7)). A logistic regression based on GAD alone, applied to the same leave-one-subject-out cross-validation method, was able to differentiate in 29/34 of the participants (85%) between first and second viewing (average accuracy 0.65±0.12). Associated AUC-ROC metrics exhibited an average of 0.68±0.15 (one-sample t-test: t(33) = 7.22, p = 2.84e^−08^). This adds to GAD’s potent predictive value and showing that distance metrics alone are sufficient for precise identification of episodic-like memory traces in the MEGA paradigm.

To further understand which eye-tracking features are most effective in capturing event memory, we incorporated SHAP (SHapley Additive exPlanations) feature importance analysis, a game theory-based method to distill and quantify each feature’s influence on the model’s output^50^. SHAP provides an average impact of each feature on model prediction by examining its performance with and without the presence of each feature across all possible feature combinations. This analysis identified distance-based features as particularly informative, with GAD ranking as the top feature (Figure 3E). Along the same lines, a waterfall plot of a representative trial illustrates how individual features (especially GAD) cumulatively influence the model’s decision-making process for a single trial, highlighting the predictive power of GAD in indexing memory in the MEGA paradigm (Figure 3F).

### Anticipatory gaze marks event recollection, whereas pupil size indexes context recognition

What aspects of memory does anticipatory gaze capture? To what degree does it reflect episodic-like memory for the event versus familiarity with the general context? To address these questions, we first tested how MEGA relates to simple verbal reports (‘Have you seen this movie before?’). We compared GAD scores in movies reported as ‘seen before’ vs. ‘not seen before’, focusing only on trials with reports associated with high confidence ratings (Figure 1B). We found a significant reduction in GAD upon 2^nd^ viewing for movies that were reported to be seen before (GAD_1st v._= 18.1 ± 0.75, GAD_2nd v._= 16.9 ± 1.1, t(33)=5.79, *p*=1.748e^−06^, Cohen’s D = 1.29, 95% CI [0.81 1.76], Figure 4A) but also a significant effect for movies that (incorrectly) reported as ‘not seen before’ (GAD_1st v._= 18.5 ± 2, GAD_2nd v._= 17.58 ± 1.85, t(33)=5.24, *p*=9.105e^−06^, Cohen’s D = 0.48, 95% CI [0.11 0.85], Figure 4A). To directly compare the difference between the first and second viewings of the explicitly recognized and not-recognized movies, we computed the normalized decrease of GAD from 1^st^ to 2^nd^ viewing (GAD^1st^ ^viewing^ - GAD^2nd^ ^viewing^ / max(GAD^1st^ ^viewing^, GAD^2nd^ ^viewing^), Figure 4B; applying other or no normalization schemes yielded similar results, Methods). Higher values reflect an anticipatory effect during the second viewing and thus reflect memory-guided behavior. This score exhibited a trend towards higher values for movies that were subsequently recognized than for not-recognized movies (M _MEGA recognized_=0.07 ± 0.07, M_MEGA not recognized_=0.05 ± 0.05, paired t-test: t(33)=1.99, *p*=0.055, Cohen’s D = 0.35, 95% CI [0.00 0.71], Figure 4B and time course in Supplementary Figure S4).

To better understand what aspects of memory are captured by anticipatory gaze beyond ‘Have you seen this movie before?’, a second experiment was conducted to evaluate multiple dimensions of memory reports in detail, aiming to distinguish between event recollection and context recognition (Figure 4C). Participants watched 48 animated movies depicting similar surprising events. After a four-hour break, they watched these movies again together with 12 novel movies in a randomized order. However, in this experiment, the 2^nd^ session included the exact same movies, except the surprising event was omitted. Because the surprising event was not presented during the 2^nd^ viewing, we could follow up with an array of retrieval tasks immediately after each movie. These tasks aimed to provide additional sensitivity to better investigate the relationship between explicit memory and anticipatory gaze. We collected the following five verbal reports: recognition, free recall, object recognition, event location recall, and temporal recall (for details see Methods).

Results reliably replicated the anticipatory gaze effect with a new group of participants. 30/30 of participants (100%) demonstrated significantly greater gaze proximity to the event location during the second viewing (M_MEGA-score_=0.086 ± 0.037, t(29)=12.8, *p*=2e^−13^, Cohen’s D = 2.34, 95% CI [1.61 3.07], Figure 4D). Next, we analyzed the GAD of each movie based on its explicit report in three categories (Methods): event recollection (correct movie recognition, as well as object recognition and event location recall), context recognition (scenery was recognized, but the object recognition or event location recall was incorrect), and unrecognized movies (incorrect recognition task, independent of the answer in event location recall or object recognition). Temporal recall (when precisely the event occurred) was not further analyzed because retrieval performance was at chance (t(29) = 0.45, p-value = 0.65). Behaviorally, retrieval performance for movie recognition, object recognition, and location recall were all significantly above chance (recognition: t(29)= 15.0, p-value = 3.21e^−15^, object recognition: t (29) = 36.8, p-value < 2.2e^−16^, location recall: t(29) = 11.8, p-value = 1.36e^−12^, Suplementary Figure S1).

Analyses revealed that MEGA scores were significantly higher for movies where participants recollected the event in full, highlighting the sensitivity of MEGA to episodic-like recall (Figure 4E & 4F). Accordingly, ANOVA with the dependent variable MEGA score and the factor memory content (event, context, and no memory) revealed a significant main effect (F(2,87)=14.11, *p*=4.9e^−06^, η² = 0.24, 95% CI [0.12, 1.00]). Post-hoc pairwise comparison revealed a significantly higher MEGA-score for full recollection of the event compared to MEGA-scores of movies where only context was recognized (M_event_=0.12 ± 0.06, M_context_=0.08 ± 0.04, *p*_Tukey_=0.002) or compared to unrecognized movies (M_event_=0.12 ± 0.06, M_no-recognition_=0.05 ± 0.05, *p*_Tukey_<0.001), but the latter two conditions did not differ significantly (context vs. no-recognition: p_Tukey_=0.2). In fact, Bayesian statistics suggest that the anticipatory gaze for movies where the scenery was familiar is similar to the ones of movies that were not recognized (BF_10_ = 0.32, prior used according to the effect size of the M_event_ vs M_no-recognition_ comparison).

Moreover, the MEGA score of each movie correlated with participants’ precision in reporting the event location such that stronger anticipatory gaze effects are associated with higher proximity to the event location (pearson *r* = 0.22, *p*<2.2e^−16^, 1438 movies). Accordingly, in each trial, the higher the precision in explicitly reporting the event location, the closer the anticipatory gaze was to that location before it occurred on the screen. Accordingly, the MEGA score of participants correlated with the number of trials categorized as event recollection (pearson *r*=0.37, *p*=0.043, N=30) but did not correlate with the number of trials that were not recognized (*r*=−0.046, *p*=0.81). Correlation with the number of trials where only the context was recognized exhibited a marginally significant negative correlation (*r*=−0.36, *p*=0.05).

Finally, we investigated anticipatory gaze within the context of participants’ free recall of surprising events, rather than location recall and forced-choice object recognition. As expected, the MEGA score was bigger if the object of a surprising event was explicitly recalled, compared to movies for which participants could not remember the event’s object (free recall: MEGA score _recalled_=0.14 ± 0.08, MEGA score _not recalled_=0.07 ± 0.03, paired t-test: t(28)=5.46, *p*=7.8e^−06^, Cohen’s D =1.22, 95% CI [0.71 1.72]). Furthermore, this difference in anticipatory gaze linearly increased for the participants’ ratio of objects recalled and forgotten (pearson *r*=0.56, *p*=0.0016).

To further distinguish MEGA from context recognition, we focused on pupil dilation as an index of familiarity upon repeated stimulus presentation^51–54^. Specifically, previous findings suggest that pupil dilation is increased for recognized words compared to unrecognized words^55,56^. In line with this literature, we also found that pupil size was larger for the second viewing compared to the first viewing (pupil size _1st v_=4436. 25 ± 378.12, pupil size _2nd v_=4647.57 ± 378.12, t = 59, p-value = 1.974e^−05^). Next, we compared changes in pupil size across repeated movie viewings in relation to the verbal report. To this end, we computed a normalized pupil size score using the same formula used for the MEGA score (1st viewing - 2nd viewing / max(1st viewing, 2nd viewing), negative values reflect a larger increase in pupil size, Methods). We found a larger increase in pupil size for movies that were recognized compared to movies that were not recognized (higher pupil size score for ‘not recognized’ in Figure 4G). ANOVA with pupil dilation score as the dependent variable and the explicit memory questions as a factor revealed a significant main effect (F(2,87)=5.05, *p*=0.008,, η² = 0.1, 95% CI [0.02, 1.00]). Post-hoc pairwise comparison revealed a significantly higher pupil score for movies where the event was recollected in full (M_event_=−0.059 ± 0.048, M_no-recognition_=−0.021 ± 0.054, *p*_Tukey_=0.013) and for movies where the context was recognized (M_context_=−0.054 ± 0.049, M_no-recognition_=−0.021 ± 0.054, *p*_Tukey_=0.033) compared to unrecognized movies. Crucially, pupil dilation score did not differ between the movies with event recollection and context recognition, suggesting no specific relationship with event memory (event vs context: p_Tukey_=0.93, BF_10_ = 0.30 ± 0.03%, default prior = 0.707). This was stable over the whole pre-event time interval (Supplementary Figure S5). These findings suggest that while the pupil is indicative of recognition of the context alone, the anticipatory gaze is guided by a richer memory that includes the recollection of event details, such as its location within the context.

### Anticipatory gaze reflects relational memory rather than familiarity

Next, we conducted another control experiment to investigate the possibility that gaze differences during second viewing might reflect familiarity with scenery, rather than relational memory. Specifically in our example movie (Supplementary Movie S2), there is the possibility that after watching the giraffe for the second time, the viewer - due to familiarity or just being bored - looks systematically away from that location and/or closer to the upcoming surprising event. Could this account for the anticipatory gaze, independent of the relational retrieval of the SE? To test this, we ran a control experiment while presenting the movies without surprising events in the first or the second viewings (i.e. just the same background scenery). A reduction of GAD (distance to SE) between first and second viewing without any events would suggest that familiarity can explain anticipatory gaze effects, while lack of GAD difference between viewings would suggest that anticipatory gaze reflects relational memory. We found that GAD was significantly smaller in this control compared to Experiment 1 (M _control_=0.006 ± 0.028, t-test: t(52) = 4.186, *p* = .0001, Cohen’s D = 1.8, 95% CI [0.6 1.8]) and Experiment 2 (t-test: t(48) = 8.247, p = 2.923e^−09^, Cohen’s D = 2.4, 95% CI [1.6 3.1]), confirming that unexpected events strongly modulate anticipatory gaze behavior (Figure 4i,j). Crucially, to test whether mere familiarity with the scenery could account for anticipatory gaze shifts, we compared GAD measures between first and second viewings in this no-surprise control condition. We observed no significant change in GAD from first to second viewing (t-test: t(49) = 0.94726, p = 0.3554, 95% CI [0.5636226 1.7956199]). We confirmed this using a Bayesian analysis, which supported the null hypothesis. When anticipation magnitudes from Experiment 2 (Prior Cohen’s d = 1.34; BF_10_ = 0.22 ± 0.09%) and Experiment 1 (Prior Cohen’s d = 0.88; BF_10_ = 0.31 ± 0.04%) were incorporated as informative priors, the Bayes Factor indicated evidence against familiarity-driven effects. Thus, these results suggest that relational memory processes, rather than familiarity with the scene, underpin the anticipatory gaze patterns observed.

### Anticipatory gaze is replicated in naturalistic movies

To what extent can anticipatory gaze be revealed using other movies, not necessarily animations compiled specifically for scientific research? We set out to test the degree to which anticipatory gaze captures episodic-like memory recall in settings that closely mimic real-world experiences, with the aim of bridging laboratory research and everyday memory. To this end, we performed a third experiment, where 32 naïve participants viewed 100 YouTube videos (Figure 5A). First, it was necessary to define the surprising event location and timing since these were not defined as a-priori as in the tailor-made animations. A group of 55 independent participants marked the spatial and temporal coordinates of the surprising event. Each movie’s event location was defined as the median (X, Y, t) of their choice of coordinates. 48 movies that exhibited a maximal level of consensus (within one standard deviation of the median time or location; see Methods) were used for subsequent analysis. Once again, analysis of GAD preceding the surprising event replicated the anticipatory gaze effect, with a completely different set of stimuli and a different group of subjects. A significant increase in gaze proximity to the event location was observed upon 2^nd^ viewing in all participants but one, robustly indexing memory without report (M_1st v._=10.96 +-0.46, M_2nd v._=10.04+-0.72, paired t-test: t(30)=10.272, p=2e^−11^, Cohen’s d = 1.52, 95% CI [0.98 2.06], Figure 5BC). Increased proximity of anticipatory gaze to the event location was evident throughout the seconds leading to the event (Figure 5B). GAD declined upon the event onset in both 1^st^ and 2^nd^ presentations, reflecting gazing towards the event once it occurred, but this drop was less steep and interestingly began already *prior* to event onset, arguably since the gaze of participants was already “drawn” towards the event location by narrative cues in naturalistic movies compared to the highly unexpected event appearance in tailor-made animations. Analyzing GAD and MEGA computed separately for recognized and unrecognized movies (according to verbal report, Figure 5A) showed that the MEGA score significantly exceeded chance-level for both remembered and forgotten movies, replicating the observation in the previous experiments (explicitly remembered trials: t(30)=10.75, p=8e^−12^ Cohen’s d = 1.93, 95% CI [1.30 2.56]; for explicitly forgotten trials: t(30)=5.71, p=3e^−6^, Cohen’s d = 1.02, 95% CI [0.57 1.48], Figure 5D). Together, the results show that the anticipatory gaze effect robustly replicates with naturalistic movies, attesting to the utility of this approach in diverse contexts, including real-life situations.

### Anticipatory gaze reveals sleep’s benefit for memory consolidation without report

We demonstrate one potential application of the MEGA paradigm in terms of sleep benefits for memory consolidation. Naïve participants were recruited for a fourth experiment, viewing naturalistic videos (identical to experiment 3) in two sessions separated by a 2h break that included either a nap opportunity for some individuals (n=19) or an equally long interval of wakefulness for other individuals (n=34). First, regardless of sleep or wake, we replicated the anticipatory gaze effect with a new group of participants. We observed significantly lower GAD reflecting anticipatory gaze (Figure 5B) in 16/19 of participants (84%), substantiated by statistical analysis (M^1st^ ^v.^=11.3±0.62, M^2nd^ ^v.^=10.12±0.87, Wilcoxon signed-rank test: V(18)=190, p=4e^−6^; rank-biserial effect size = 0.88, 95% CI [0.88 0.88]). This constitutes a third successful replication of the anticipatory gaze effect, reinforcing its reliability in capturing memory. Next, comparing sleep and wake, we found that the MEGA score in the nap condition was 23.6% higher than in the awake condition (M_wake_=0.07 ± 0.048, M_nap_=0.09 ± 0.048, Wilcoxon signed-rank test: *z*(N₁ = 19, N₂ = 34) = 1.71, p=0.0439, Cohen’s d=0.48, 95% CI [0.1 1.1]). In contrast, the verbal reports did not reflect this pattern. Bayesian statistics provide evidence that explicit recognition rates were comparable following sleep and wake consolidation (BF_10_ = 0.39 ± 0%, prior used according to the effect size of the anticipation effect). Participants recognized 76.8 ± 7.1% of the movies after sleep and 74.3 ± 7.5% after wakefulness (Wilcoxon signed-rank test: W = 401, p = 0.098). This discrepancy highlights the sensitivity of the MEGA paradigm that may be masked when only considering explicit verbal reports. Overall, these results indicate that the nap had a positive impact on memory consolidation as reflected in anticipatory gaze, demonstrating the potential of MEGA as a no-report paradigm for studying the relation between sleep and episodic-like memory consolidation.

## Discussion

This study shows that tracking gaze during repeated viewings of movies with surprising events constitutes an effective method for investigating memory without verbal reports. The results establish that during the second movie viewing, the gaze gravitates towards the event location, exhibiting memory-guided prediction that anticipates its occurrence. Gaze distance (GAD) can be used as an intuitive metric to capture the degree of this predictive anticipation, showing significantly higher proximity to the event location during the second viewing. The anticipatory gaze effect was captured before changes were visible in the movie and even when the surprising event was entirely absent in the second movie viewing. This establishes that it corresponds to memory-guided prediction irrespective of the visual cues that mark it. Anticipatory gaze is a highly robust effect that is consistently observed and replicated several times across multiple stimulus types and in different naïve groups of participants (N=126) - attesting to its versatility and utility across settings. Machine learning classifications of features extracted from gaze data identify memory traces at a single-trial level in new participants. In a separate experiment where we collected verbal reports about the movie events, we found that anticipatory gaze effects are largest when the surprising event was fully recalled, showing dissociation from pupil size measures not associated with recalling the surprising event. Finally, we illustrate how applying MEGA without verbal reports can effectively replicate the classical beneficial effect of sleep on declarative memory^44,57,58^. MEGA can be successfully employed using either naturalistic movies or custom animations specifically designed for this purpose, a resource made available for any future follow-up study (https://yuvalnirlab.com/resources/).

The MEGA paradigm approach creates one-shot encoding of movies with scene-event pairs that are recalled after 2 hours. Anticipatory gaze reflects the recollection of specific events, positioning MEGA as a no-report variation of a cued recall task. By creating our own paradigm including tailor-made animations we extend previous research by Kano & Hirata^13^ on great apes, as well as adaptations of their paradigm ^36,59^. We combine elements from decade-long research on visual exploration in images^25,27–30,54^, visual search^15,16^ and contextual cueing^21,22,60^ in picture scenes, as well as long-term memory guided anticipatory spatial orienting of attention^23,61^ and anticipatory viewing behaviors ^39–43^. We integrated these elements into tailor-made animations designed to elicit memory-guided anticipatory gaze toward the location of the upcoming event, thereby maximizing single-trial accuracy of event recollection. Thus, our work adds to the growing literature on utilizing anticipatory gaze as a marker of relational memory^26,27,62^.

Two complementary approaches were used to quantify the anticipatory gaze effect. The first and intuitive metric, GAD, captures the average Euclidean distance to the location of the surprising event from the beginning of the movie presentation until the event occurs and enables identification of memory-guided gaze in ∼90% of participants. A second machine learning approach using the XGBoost classification algorithm is aimed at the predictive power of single-trial gaze. Surprisingly, machine learning applied on merely seven-second intervals of gaze data during pre-event movie viewing - without any averaging across movies or participants - was sufficient to significantly identify whether this viewing was associated with memory for the event (first or second viewing). Strikingly, classification performance was maintained even when we only used gaze distance instead of a comprehensive set of 87 gaze distance-related eye-tracking features. A post-hoc comparison of all features’ importance confirmed that GAD was most informative.

We created MEGA to capture the memory of the ‘what, where and when’ of the surprising event. But does anticipatory gaze <u>exclusively</u> reflect episodic-like memory? Although the current dataset cannot conclusively establish that anticipatory gaze exclusively reflects episodic-like memory, several reasons suggest that this is the case. First, anticipatory gaze correlated with the event recollection, its free recall, the location recall and the object recall, therefore reflecting the recollection of the SE. Even on a single-trial level, anticipatory gaze correlated with the accuracy of the participants estimation of the event location. Second, one-shot encoding of detailed long-term memory makes it unlikely to be explained by a non-declarative memory system, especially since some of the cues (scenes) are highly similar for different events. Third, we demonstrate that the anticipatory gaze vanishes if there is only familiarity of the scene (control experiment). Also, statistical learning seems unlikely because GAD during the first viewing was not lower at the end of the first (encoding) viewing session. Nonetheless, we acknowledge that robust (but reduced) anticipatory gaze was also observed for movies that were not consciously recognized. A more definitive answer regarding selectivity of our task for episodic memory may be provided with time by neuroimaging and/or lesion studies assessing the role of medial temporal lobe systems in MEGA.

What can be gained by estimating memory with MEGA, independent of verbal reports? Foremost, our new analytical approach demonstrates sufficient sensitivity to detect memory traces at the single-trial level. Specifically, our metric is correlated with the accuracy of the recollection of the event’s location, on a movie-level. This validates MEGA’s potential as a no-report paradigm for episodic-like memories Furthermore, MEGA provides an additional method and approach in clinical settings where verbal communication is challenging. For instance, individuals with cognitive impairments, such as aphasia or developmental disorders, often struggle with complex instructions or language comprehension and production. Along the same lines, a stroke patient may be unable to speak, yet we would like to estimate their memory capacity. Finally, MEGA can enhance consistency in memory research by increasing generalizability across participants who speak different languages. Finally, from a basic science perspective, current research employing reports confounds memory itself with the ability to articulate it. In this context, MEGA can help distill the brain activities and diseases affecting memory per se beyond its access and report.

Another important advantage of MEGA is that it goes beyond a binary ‘recognized’ vs. ‘non-recognized’ report to represent memory as a continuous quantitative variable. The MEGA-score metric is sensitive to various anticipatory behaviors, whether through a few prolonged fixations at the event location, multiple fixations around it, or even subtle proximity to the event location. This granularity allows us to reveal a linear relation between the anticipatory gaze scores and the individual precision in reporting the event’s location for single movies, suggesting that anticipatory gaze captures “more” than just the recognized – non-recognized distinction. Although the aggregation over the pre-event interval proved robust, the temporal dynamics of the anticipatory gaze remain inconclusive and warrant further investigation (see Figures S5 and S6).

### Limitations

At present, it is still unknown the extent to which MEGA depends on activity in the hippocampus and the medial temporal lobe (MTL). Are the neuronal underpinnings comparable to those associated with episodic memory, and/or to other implicit eye-movement-based memory effects?^51,63,64^ Additionally, it should be acknowledged that the current results involve a task where participants were required to report if the movies were previously seen or not, thereby introducing an additional layer of cognitive processing that could influence the natural recall of events. Future studies should assess how anticipatory gaze unfolds in entirely passive viewing conditions without any instructions. The present findings already extend the literature on sleep consolidation, suggesting that the benefits of sleep for memory consolidation extend beyond verbally recalling memories. However, the direct relationship between memory consolidation, as captured by MEGA, and brain activities supporting this process —such as slow waves and sleep spindles—remains unclear and require further investigation.”

Looking ahead, eye-tracking may offer a promising avenue for understanding and diagnosing memory disorders^59,59,65^. In clinical settings, our findings in healthy adults hold promise for improving and refining existing non-verbal tasks^17,66–68^ for early diagnosis of memory disorders in mild cognitive impairment (MCI) and for monitoring Alzheimer’s disease progression.

Standard cognitive and neuropsychological assessments like the MMSE^69^ or the MoCA^70^ are limited in detecting preclinical memory deficits. Beyond degeneration, our approach may provide a new perspective on how to assess memory upon damage to the medial temporal lobe (MTL)^71^, when it is often unclear to what extent damage affects mnemonic systems or their interface with other brain systems that enable conscious report. Another clinical application pertains to patients suffering from motor or language disorders that limit verbal report, such as those with aphasia^72,73^, offering a non-verbal means of assessing memory integrity.

In conclusion, the MEGA paradigm offers a valuable analytical approach and introduces a significant advance in memory research. Utilizing eye tracking to study memory without verbal reports offers wide implications for both basic research in cognitive neuroscience and clinical fields.

## Acknowledgments

We extend our deepest gratitude to Dr. Noa Regev for her administrative assistance and unwavering support throughout this project. We are indebted to Yuval Shapira for their initial analyses and for developing a user interface that enabled participants to accurately select event coordinates in natural movies. Special thanks go to Odessa Goldberg, Alexandra Klein, Yael Gat, Rotem Falach, Or Ra’anan, and the additional research assistants for their invaluable help with data collection. We also want to express our heartfelt thanks to the participants of this study, whose involvement was crucial to the success of our research. Additionally, we are grateful to Studio Plonter® for their exceptional work in creating the tailor-made animations that played a pivotal role in our experiments. We are also thankful to Dr. Roy Amit for his advice on eye-tracking data collection and preprocessing in the initial stages of the project.

This research was supported by ERC-2019-CoG 864353 (Y.N.). Additionally, F. J. Schmidig was supported by a Tel Aviv University Sagol School of Neuroscience Postdoctoral Fellowship. The funders had no role in study design, data collection and analysis, decision to publish or preparation of the manuscript.

## Author contributions

- Conception and Design of Research: D. Yamin, O. Sharon, F.J. Schmidig, and Y. Nir.
- Funding Acquisition: Y. Nir and O. Sharon.
- Compilation of Movie Stimuli: O. Sharon and D. Yamin.
- Data Collection: D. Yamin, F.J. Schmidig, O. Sharon, and Y. Nadu.
- Data Analysis: D. Yamin, F.J. Schmidig, O. Sharon, and J. Nir.
- Manuscript Writing: D. Yamin, F.J. Schmidig, and Y. Nir.
- Advice on Experimental Design and Data Analysis: C. Ranganath.
- Critical Review of Results and Manuscript Comments: All authors.

## Data availability

Source data underlying the results presented in figures can be found here^74^: https://osf.io/b64qk/

Additional data will be provided upon reasonable request to the corresponding author.

## Code availability

The code used to analyze the gaze data in this study is available at: https://github.com/dyamin/MEGA and in https://osf.io/b64qk/

Specific code for detailed analysis is available from the authors upon reasonable request. The experiment code is available at: https://github.com/dyamin/MEGA-Experiment

The stimuli used in the experiments are publicly available at: https://yuvalnirlab.com/resources/

