## Supplementary information for "Anticipatory Eye Gaze as a Marker of Memory"

### Supplementary Figure 1: Anticipatory Gaze and Explicit Memory Reports

In Experiment 2, anticipatory gaze was evaluated in relation to event-specific recollection. Based on four forced-choice tasks (i - recognition, iii - what, iv - where, v - when), we defined three labels of remembered memory content: (A) Context and event recollection: movies where participants accurately recall the context and the surprising event. Namely, a correct answer for the movie recognition, object recognition, and the event location recall. (B) Context recognition: recognition of the movie's setting without specific recollection of the event. Namely, a correct answer for the movie recognition but an incorrect answer for the object recognition and location recall, and (C) not recognized: when participants did not recognize the movie at all. Please note that location accuracy was defined as correctly identifying the quadrant of the upcoming event. the Temporal recall was not further analyzed since retrieval performance was at chance (t(29) = 0.45, p-value = 0.65). The free recall (ii) was analyzed separately. Here in figure S1 we display the raw responses to each separate question.

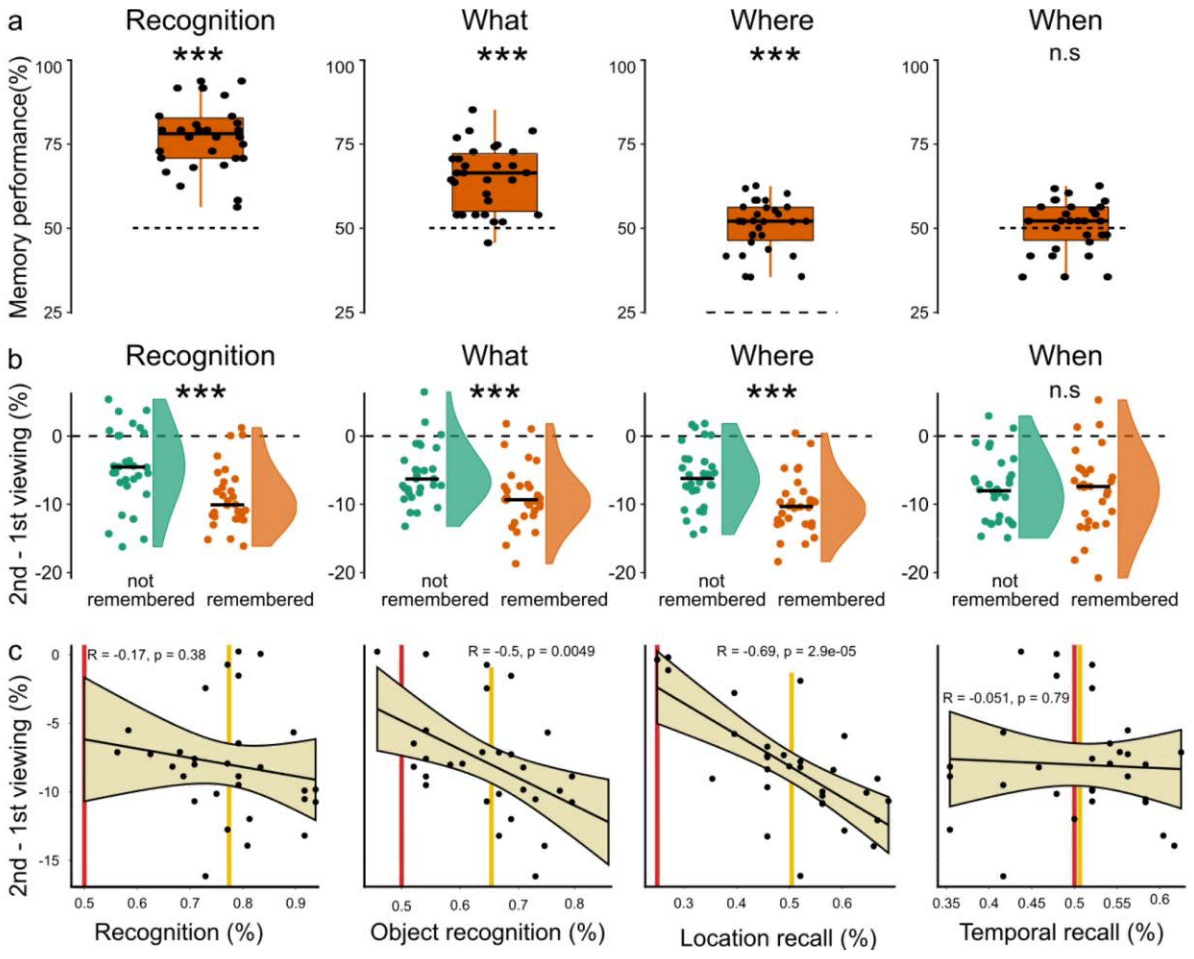


**Figure S1. The relationship between anticipatory gaze and explicit memory reports – original retrieval questions** A) verbal report: Participants' retention rate for each retrieval task independently. Tasks from left to right: movie recognition, object recognition, spatial recall, temporal recall. B) Memory task and MEGA-score: Comparison between anticipatory gaze of correct (orange) vs incorrect (green) answers, according to each retrieval task. Anticipatory gaze is higher for movies that were recognized and higher if participants recalled the object and the location of the SE. C) Correlation of individuals' anticipatory gaze score and their verbal reports, indicating more object recognition and higher spatial recall relates to the MEGA-score. * = p<0.05, ** = p<0,01, *** = p<0,001, dotted lines mark chance-level performance.

### Supplementary Figure 2: Exemplary Viewing Behaviour

**Movie S1. Exemplary viewing behaviour during an animated movie and naturalistic movie.** We present the viewing behaviour of one participant while watching an animation or a naturalistic movie. The download link points to four videos consists of 2 movies used in the first two experiments and 2 movies used in Experiment 3. Superimposed on the movie is the gaze of a particular subject. The blue dots represent gaze during the first viewing and green dots mark the gaze during the second viewing. The red dot marks the center of the upcoming surprising event. In addition, on the top left we display the distance of the momentarily gaze to the location of the upcoming surprising event. The difference between the two distances represent anticipatory gaze (before standardization).

### Supplementary Figure 3: Stimuli

**Animation Movies (Experiments 1 & 2).** For Experiment 1, we used 65 silent animated colored movie clips (see examples in Supp. Movies A1 and A2), each rendered at a resolution of 2304x1296 pixels. These clips varied in duration from 8 to 22 seconds, with an average length of 12.68 seconds. Movies were prepared with the aim of being universally comprehensible, regardless of age, cultural background, or cognitive ability. A key feature of these animations was the inclusion of a surprising event designed to be unmistakably noticeable upon first viewing while adhering to the following criteria: i) The event occurred after a minimum of 5 seconds from the start of the clip, ensuring viewers are engaged and have enough time to potentially recognize the context. ii) Once introduced, the event remained visible for at least 2 seconds. iii) It appeared peripherally, not centrally, varying in location and timing across clips to prevent predictability. iv) There were no hints to spoil the surprise and indicate the event's appearance. Additionally, the movies were crafted to include background scenes that are visually similar, yet not identical, across different movie clips (e.g., more than one movie had the same underground water scenery) to increase difficulty and avoid ceiling performance. Animations were crafted by Studio Plonter® (https://www.plonteranimation.com/). For the second viewing in Experiment 2, the animation studio produced an alternative version of 60 animations, identical in every aspect but without the surprising event. Visual stimuli were presented using a 15.6-inch (35x20 cm) monitor with a resolution of 3840x2160 pixels.

**Example scenery and objects:**

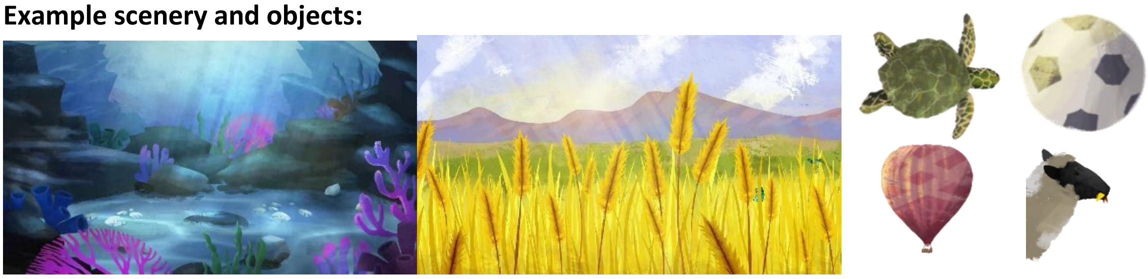

**Naturalistic movies (Experiments 3 & 4).** Movies (examples in Supp. Movies R1 and R2) included 100 silent black and white movie clips with a duration of 3.5 to 26 seconds downloaded from YouTube. This collection encompassed carefully selected clips from classic films (e.g., Charlie Chaplin) or contemporary single-shot natural scenes characterized by minimal camera movement, containing at least one SE. Scenes covered a broad spectrum of topics ranging from famous soccer goals through animal behaviors to human social interactions. Visual stimuli were presented using a 24-inch monitor (51x29 cm) with a resolution of 1920x1080 pixels.

**Determining the surprising event.** Determining the surprising event is central for quantification to the anticipatory gaze, as event onset needs to be defined for data analysis. For animated movies (Experiments 1, 2), the animation studio defined the location and timing of the surprising events. The onset and location of the surprising event were defined by the first pixel of the event, which becomes visible on the screen. For naturalistic movies (Experiments 3,4) the surprising event was defined experimentally by an online human focus group. We followed protocols used in prior research[72–74] and conducted an additional dedicated online experiment with 55 participants who were asked to mark the time and location of the surprising event E in a random subset of the 80 movies used for analysis (excluding the 20 unseen and those that were replaced by them, M = 51.05±1.16 rated movies per participant). Participants watched the movies and then, during an additional viewing, pressed their mouse at the moment (t) and location (x, y coordinate) where the surprising event occurred. Each movie's event onset, duration, and location were then defined as the median of marked events. We discarded from the study movies that did not reach a minimal consensus on the timing and location of the surprising event since such movies typically contained multiple events and could not yield consistent eye-tracking patterns. Accordingly, movies where less than 70% of participants agreed on the location were excluded (consent meant localizing the location within 1/9 of the screen size). Overall, 48 of the 80 YouTube movies (60%) were deemed suitable for eye-tracking experiments.

### Supplementary Figure 4: Time Course of Anticipatory Gaze

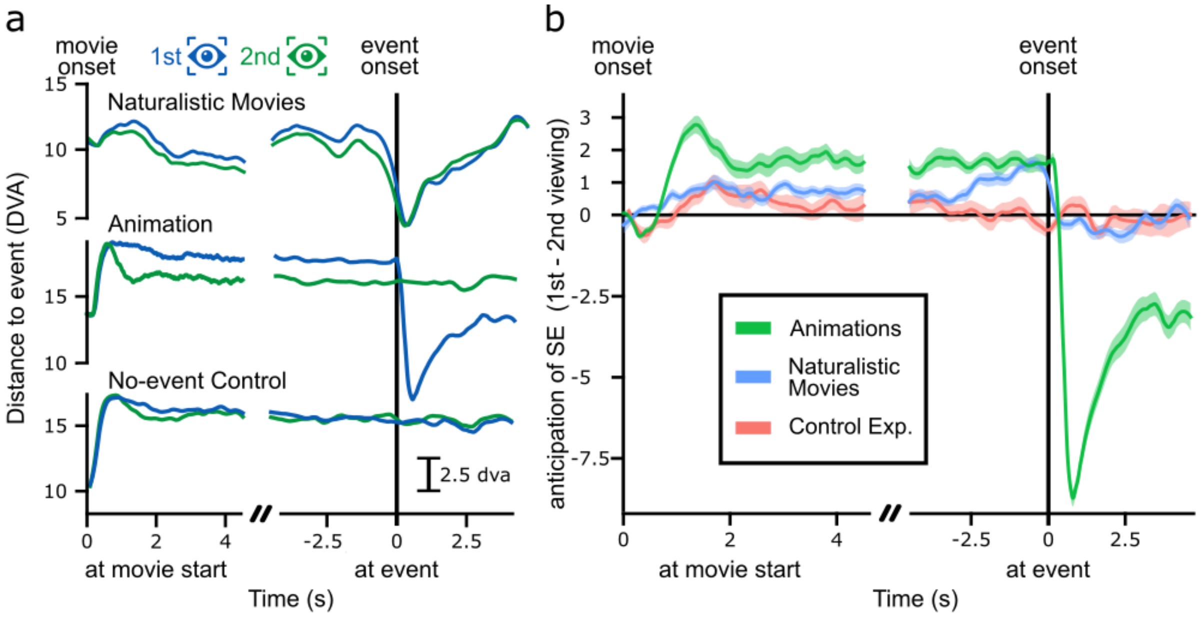


**Anticipatory gaze related to event boundary.** To examine the time course of the anticipatory gaze we aligned the gaze to the movie onset (left) and to the surprising event (right) and cropped both images together. A) Comparison of the average distance of the gaze to the location of the SE between the first and second viewing for experiment 3 (naturalistic set), experiment 2 (animation set) and the control experiment. Please note that in experiment 2 the surprising event was removed such that an explicit report could take place. B) Direct comparison of the differences in viewing behaviour between the two viewings in A, reflecting anticipation of the SE. Shaded areas demonstrate variance between subjects as standard error of the mean.

The temporal dynamics differ between real-life videos and animations, particularly in how viewers anticipate surprising events. In real-life clips, viewers often anticipated the surprise even already during the first viewing, influenced by contextual cues such as actors' gaze or body orientation, or camera/zoom motion, leading to a gradual drop in the GAD (gaze anticipation distance) before the event. Nevertheless, anticipation increased towards the events' timing (Figure S5b). In contrast, animations were designed to avoid such cues, resulting in no early GAD drop, but also making it harder to predict when surprises occur. Across both types, a consistent GAD difference emerges within the first second, possibly reflecting the recognition of the scene and the recollection of the SE, including its location.

### Supplementary Figure 5: Time Course of Pupil Dilation

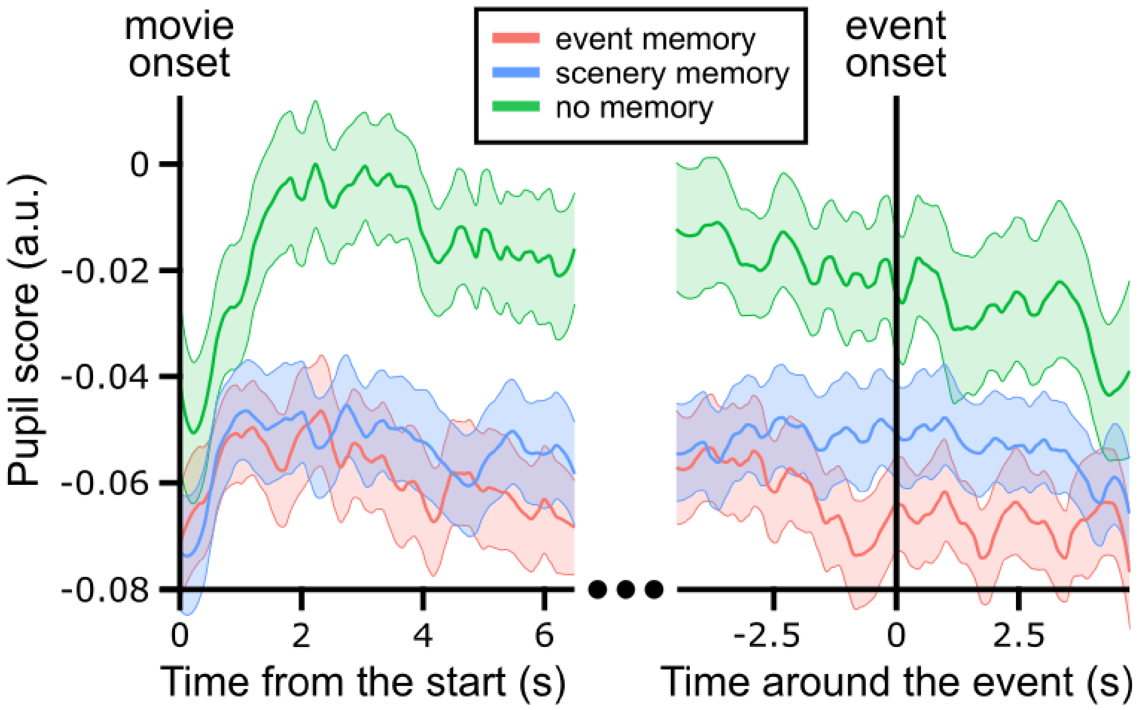


**Pupil dilation related to event boundary.** To visualize the temporal dynamics, we aligned the pupil dilation to the movie onset (left) and to the surprising event (right). This way, we average over all subject and movies for each timepoint in relation to the crucial event boundary (once onset & once SE). At the end we cropped both figures together, such that the right side of the figure is not a continuation of the left side. Each line depicts the normalized score of the pupil dilation (as the MEGA score, see methods). The normalized pupil score is based on the difference between the pupil dilation of both viewings. Please note that the SE was cropped out in the second viewing, where they were present during the first viewing. To examine the difference of the pupil dilation in respect of the subsequent memory effect, we compare movies when participant recollected the SE (red), participants recognized the scenery but did not recall the event (blue) and where the movie was wrongly identified as new. Bold line reflects the average of all participants pupil dilation and shaded areas depict standard error of the mean between subjects.

From the temporal dynamics we infer that the difference between movies without a subsequent memory effect (“no memory”) and with a subsequent memory effect (“event memory” & “scenery memory”) emerges within the first two seconds after movie onset. Furthermore, these results support our finding that the pupil dilation does not differentiate between event memory and scenery memory, at least they do not invalidate them.

### Supplementary Notes 6: Single-Trial Decoding of Movie Viewing

**Feature engineering for machine learning (ML) analysis.** Our ML analysis relied on extracting and transforming raw eye-tracking data into a set of features. We calculated a total of 243 eye-tracking features. These features were broadly categorized into four groups: Fixation, Saccades, Blinks, and Pupil. Each category encapsulated different aspects of the participants' eye movements and responses. The features were primarily computed using each movie clip's time interval leading up to the event onset. Of these 243 features, 87 were related to the event: where the event was about to appear ("Event-Based Features"). These features leveraged the location and timing of the event, offering a more nuanced view of how participants' gaze interacted with specific elements of the visual stimuli. A comprehensive list and description of the Event-Based Features can be found in Supp. Table T4.

**XGBoost Classifiers.** ML employed an ensemble of XGBoost (Extreme Gradient Boosting) classifiers, operating through the construction of multiple decision trees in a gradient boosting framework, where each tree is sequentially built to correct the errors of its predecessors. The optimization of node splits and tree structure is driven by gradient statistics of a loss function, aiming to minimize prediction errors. Our implementation adhered to the standard specifications of XGBoost, as delineated by Chen and Guestrin[40]. The algorithm was implemented by its application to each fold in our leave-one-subject-out (LOSO) cross-validation (CV) framework, ensuring robust and generalizable findings. LOSO iteratively trains the model on the dataset excluding data from one subject, which is then used as the test set. This approach mitigates potential biases and accounts for inter-individual variability, enhancing the model's applicability to new, unseen data. To optimize parameters, we used a grid search process to pinpoint optimal hyper-parameters within predefined ranges: 'learning_rate' (0.001-0.5), 'max_depth' (2-4), and 'n_estimators'(100-150). Final average hyperparameters used were 'learning_rate': 0.058, 'max_depth': 3.26, and 'n_estimators': 118.52, were instrumental in fine-tuning our models to achieve optimal performance.

**Performance evaluation.** We quantified model performance via several metrics: A) Classification Accuracy: the proportion of correctly predicted viewings, capturing the model's ability to accurately differentiate between 1$^{st}$ and 2$^{nd}$ viewings. B): Average Confusion Matrix: a more nuanced breakdown of model predictions separating true positives, true negatives, false positives, and false negatives. C) Receiver Operating Characteristic (ROC) Curve: The ROC curve shows the trade-off between the true positive rate and false positive rate across different thresholds, with the Area Under the Curve (AUC-ROC) providing a summary measure of model performance. The statistical significance of ML classification was quantified by applying a one-sample t-test to the ROC AUC score compared to the chance level (50%).

**SHAP Analysis.** Understanding the intricate decision-making processes of our ensemble, XGBoost classifiers necessitated the integration of SHAP (SHapley Additive exPlanations) analysis grounded in cooperative game theory[41], to interpret the contribution of each feature to individual predictions. SHAP analysis was implemented in Python 3.9 Anaconda, with the shap package version 0.42.1.

### Supplementary Notes 7: Features used for Single-trial Classification

**Table 1: Event-Based Eye-Tracking Features.** An exemplary list of features used for the XGBoost classification of first and second movie viewings. All features are in relation to the surprising event. *SEC*: surprising event center point.

| **Feature** | **Description** |
|---|---|
| **Raw Gaze** | |
| In SEC Gaze Count | Total number of gaze points within the SEC. |
| Out SEC Gaze Count | Total number of gaze points outside the SEC. |
| In/Out Gaze Ratio | The ratio of in SEC data points to out SEC data points. |
| Gaze Entry Count | The number of entries the gaze made into the SEC. |
| Gaze Entry Rate | Frequency of gaze entries into the SEC per unit of time. |
| Mean Euclidean Distance to SEC | Average Euclidean distance from gaze points to the center of the SEC. |
| Median Euclidean Distance to SEC | Median Euclidean distance from gaze points to the center of the SEC. |
| Minimum Euclidean Distance to SEC | Smallest Euclidean distance from any gaze point to the SEC. |
| Maximum Euclidean Distance to SEC | Largest Euclidean distance from any gaze point to the SEC. |
| STD of Euclidean Distance to SEC | Variability of Euclidean distances from gaze points to the SEC. |
| SEM of Euclidean Distance to SEC | Standard error of the mean of Euclidean distances to the SEC. |
| AUC for Euclidean Distance to SEC | Area under the curve for Euclidean distance measurements to SEC. |
| **Fixation** | |
| SEC Fixation Count | Total number of fixations within the SEC. |
| Out SEC Fixation Count | Total number of fixations outside the SEC. |
| In/Out Fixation Ratio | Ratio of in SEC fixations to out SEC fixations. |
| Fixation Entry Count | Number of times fixations entered the SEC. |
| Fixation Entry Rate | Frequency of fixation entries into the SEC per unit of time. |

| | |
|---|---|
| Mean Fixation Duration | Average duration of fixations within the SEC. |
| Median Fixation Duration | Median duration of fixations within the SEC. |
| Max Fixation Duration | Longest single fixation within the SEC. |
| Min Fixation Duration | hortest single fixation within the SEC. |
| STD of Fixation Duration | Variability in the duration of fixations within the SEC. |
| SEM of Fixation Duration | Standard error of the mean of fixation durations within the SEC. |
| AUC for Fixation Duration | Area under the curve for fixation durations within the SEC. |
| **Saccade** | |
| Saccade-start SEC Count | Number of saccades that started within the SEC. |
| Out SEC Saccade-start Count | Total number of saccades that started outside the SEC. |
| In/Out Saccade-start Ratio | Ratio of in SEC saccade-starts to out SEC saccade-starts. |
| Saccade-end SEC Count | Number of saccades that ended within the SEC. |
| Out SEC Saccade-end Count | Total number of saccades that ended outside the SEC. |
| In/Out Saccade-end Ratio | Ratio of in SEC saccade-ends to out SEC saccade-ends. |
| Saccades to First Fixation within SEC | Number of saccades before the gaze first fixates within the SEC. |
| Mean Saccade Peak Velocity to SEC | Average Velocity towards the SEC |
| Min Saccade Peak Velocity to SEC | Minimum peak velocity of saccades moving towards the SEC. |
| Max Saccade Peak Velocity to SEC | Maximum peak velocity of saccades moving towards the SEC. |
| STD of Saccade Peak Velocity to SEC | Variability in the peak velocity of saccades towards the SEC. |
| SEM of Saccade Peak Velocity to SEC | Standard error of the mean of saccade peak velocities towards the SEC. |
| AUC for Saccade Peak Velocity to SEC | Area under the curve for saccade peak velocities towards the SEC. |
| **Visual Angle** | |
| Mean Visual Angle from SEC | Average visual angle from the SEC. |

| | |
|---|---|
| Minimum Visual Angle from SEC | Smallest visual angle from the SEC. |
| Maximum Visual Angle from SEC | Largest visual angle from the SEC. |
| STD of Visual Angle from SEC | Variability in the visual angle from the SEC. |
| SEM of Visual Angle from SEC | Standard error of the mean of visual angle measurements from the SEC. |
| AUC for Visual Angle from SEC | Area under the curve for visual angle measurements from the SEC. |
| Mean Visual Angle to SEC | Average visual angle to the SEC. |
| Minimum Visual Angle to SEC | Smallest visual angle to the SEC. |
| Maximum Visual Angle to SEC | Largest visual angle to the SEC. |
| STD of Visual Angle to SEC | Variability in the visual angle to the SEC. |
| SEM of Visual Angle to SEC | Standard error of the mean of visual angle measurements to the SEC. |
| AUC for Visual Angle to SEC | Area under the curve for visual angle measurements to the SEC. |
| **Pupil during looking at SEC** | |
| Minimum Pupil Radius | Minimum pupil size difference when looking at versus away from the SEC |
| Maximum Pupil Radius | Maximum pupil size difference when looking at versus away from the SEC |
| Average Pupil Radius | Average pupil size difference when looking at versus away from the SEC |
| Median Pupil Radius | Median pupil size difference when looking at versus away from the SEC |
| STD Pupil Radius | STD of pupil size difference when looking at versus away from the SEC |
| SEM Pupil Radius | SEM of pupil size difference when looking at versus away from the SEC |
| AUC Pupil Radius | AUC of pupil size difference when looking at versus away from the SEC |
| Pupil Radius first glance | Pupil radius of first fixation inside minus the previous outside the SEC. |
